# Tropomyosin Tpm3.1 is required to maintain the structure and function of the axon initial segment

**DOI:** 10.1101/711614

**Authors:** Amr Abouelezz, Holly Stefen, Mikael Segerstråle, David Micinski, Rimante Minkeviciene, Edna C. Hardeman, Peter W. Gunning, Casper C. Hoogenraad, Tomi Taira, Thomas Fath, Pirta Hotulainen

## Abstract

The axon initial segment (AIS) is the site of action potential initiation and serves as a vesicular filter and diffusion barrier that help maintain neuronal polarity. Recent studies have revealed details about a specialized structural complex in the AIS. While an intact actin cytoskeleton is required for AIS formation, pharmacological disruption of actin polymerization compromises the AIS vesicle filter but does not affect overall AIS structure. In this study, we found that the tropomyosin isoform Tpm3.1 decorates a population of relatively stable actin filaments in the AIS. Inhibiting Tpm3.1 in cultured hippocampal neurons led to the loss of AIS structure, the AIS vesicle filter, the clustering of sodium ion channels, and reduced firing frequency. We propose that Tpm3.1-decorated actin filaments form a stable actin filament network under the AIS membrane which provides a scaffold for membrane organization and AIS proteins.

## INTRODUCTION

The proximal ends of axons in the vertebrate nervous system contain the axon initial segment (AIS). The AIS serves as the site of action potential initiation and plays a role in maintaining neuronal polarity. The clustering of sodium channels at the AIS facilitates spike generation (Kole et al., 2008), while its role in maintaining polarity is the result of a vesicle filter and diffusion barrier that restrict the entry of dendritic proteins and membrane lipids into the axon (Brachet et al., 2010; Nakada et al., 2003; Song et al., 2009; Sun et al., 2014; Winckler et al., 1999). The AIS is a remarkably stable structure comprising a specialized membrane and protein complex. Central to this complex is ankyrin G (Kordeli et al., 1995; Rasband, 2010), which acts as an adaptor that recruits other AIS proteins (Jenkins and Bennett, 2001); ankyrin G recruits and binds to βIV-spectrin (Yang et al., 2007), neurofascin-186 (NF-186) (Ango et al., 2004), as well as sodium (Zhou et al., 1998) and KCNQ2/3 channels (Pan et al., 2006). The loss of ankyrin G leads to the loss of all other AIS components (Hedstrom et al., 2008; Jenkins and Bennett, 2001; Zhou et al., 1998). The interaction of ankyrin G with microtubules (Freal et al., 2016; Kuijpers et al., 2016; Leterrier et al., 2011) and the binding of βIV-spectrin to actin filaments (Jenkins and Bennett, 2001; Leterrier et al., 2015) link the AIS complex to the cytoskeleton.

While recent studies have shed light on the role of microtubules in establishing and maintaining the AIS (Freal et al., 2016; Klinman et al., 2017; Kuijpers et al., 2016; van Beuningen et al., 2015), the precise role of actin in the AIS remains unclear (Papandreou and Leterrier, 2018). Proper AIS development requires an intact actin cytoskeleton (Xu and Shrager, 2005), but the mature AIS is insensitive to actin-disrupting drugs (Abouelezz et al., 2019; Jones et al., 2014; Leterrier et al., 2015; Qu et al., 2017; Sanchez-Ponce et al., 2011; Song et al., 2009). This suggests that actin has no role in maintaining the structure of the AIS. Alternatively, actin filaments in the AIS may be resistant to the action of actin-disrupting drugs due to a low rate of turnover. Nonetheless, the integrity of the actin cytoskeleton is important for the AIS vesicle filter and diffusion barrier (Al-Bassam et al., 2012; Nakada et al., 2003; Song et al., 2009; Winckler et al., 1999). Platinum replica electron microscopy showed that the AIS contains both short, stable actin filaments, as well as longer, dynamic filaments (Jones et al., 2014). Actin-based myosin motors play a role in the targeted delivery of somatodendritic and axonal vesicles (Janssen et al., 2017; Lewis et al., 2011; Lewis et al., 2009), and actin filaments form patches in the AIS (Balasanyan et al., 2017; Watanabe et al., 2012) that may serve as vesicle filters (Janssen et al., 2017; Leterrier and Dargent, 2014; Watanabe et al., 2012). While actin patches are not exclusive to the AIS, AIS actin patches are more stable than patches along the distal axon; the washing away of diffuse molecules by cell permeabilization led to the loss of actin patches in the distal axon, but not in the AIS (Watanabe et al., 2012). In addition, recent work showed that the actin-based non-muscle myosin II is involved in AIS structure and plasticity (Berger et al., 2018; Evans et al., 2017) and associates with axonal F-actin (Wang et al., 2018).

Super-resolution microscopy revealed the presence of periodic, sub-membranous, adducin-capped actin rings in the axon, forming a lattice with spectrin and ankyrin (D’Este et al., 2015; Leite et al., 2016; Leite and Sousa, 2016; Leterrier et al., 2015; Xu et al., 2013; Zhong et al., 2014). This structure bears a striking resemblance to the erythrocyte membrane skeleton (Bennett and Baines, 2001; Leite et al., 2016; Leite and Sousa, 2016; Xu et al., 2013), where short, stable, adducin-capped actin filaments also form a sub-membranous lattice with spectrin and ankyrin (Fowler, 2013). These filaments are also partly stabilized by tropomyosins (Fowler and Bennett, 1984; Sung et al., 2000; Sung and Lin, 1994). Tropomyosins are actin-binding proteins that form coiled-coil dimers and play a role in the regulation of the actin cytoskeleton in an isoform-specific manner (Gunning et al., 2008). Of the two tropomyosin isoforms in the erythrocyte membrane skeleton, only isoform Tpm3.1 is found in the brain (Schevzov et al., 2005b). Tpm3.1 localizes to the axons of developing neurons and was suggested to play a role in neuronal polarity (Hannan et al., 1995; Vindin and Gunning, 2013; Weinberger et al., 1996). In addition, Tpm3.1 plays a role in regulating the filamentous actin pool in growth cones (Schevzov et al., 2008), growth cone motility (Fath et al., 2010), and neurite branching (Schevzov et al., 2005a). Tpm3.1 binds F-actin with high affinity (Gateva et al., 2017) and regulates the actions of key actin-binding proteins: (i) Tpm3.1 inhibits Arp2/3 complex-mediated polymerization (Kis-Bicskei et al., 2013), (ii) Tpm3.1 enhances the phosphorylation of actin-depolymerizing factor/cofilin (Bryce et al., 2003), thus inhibiting filament severing as well as depolymerization at the pointed ends (Broschat, 1990), (iii) Tpm3.1 recruits tropomodulin to the pointed ends (Sung and Lin, 1994), further lowering the rate of depolymerization (Weber et al., 1994; Yamashiro et al., 2014), and (iv) Tpm3.1 recruits and activates myosin II (Bryce et al., 2003; Gateva et al., 2017). In this study, we show that Tpm3.1 is part of the AIS actin cytoskeleton and is necessary for the maintenance of AIS structure and function.

## RESULTS

### Tropomyosin isoform Tpm3.1 is part of the AIS actin cytoskeleton

Actin filaments in the AIS organize into periodic, sub-membranous, adducin-capped rings, forming a lattice with spectrin and ankyrin (D’Este et al., 2015; Leite et al., 2016; Leite and Sousa, 2016; Leterrier et al., 2015; Xu et al., 2013; Zhong et al., 2014). We used structured illumination microscopy (SIM) to test if Tpm3.1 also decorated actin filaments in these sub-membranous rings in cultured rat hippocampal neurons at 14 days-*in-vitro* (DIV). We used anti-γ/9d and Alexa 488-tagged phalloidin to visualize Tpm3.1 and F-actin, respectively. Anti-ankyrin G served to label the AIS. We optimized imaging parameters to visualize anti-γ/9d, tagged using Alexa-647. Similar to sub-membranous F-actin, Tpm3.1 showed a periodic distribution in the AIS (Fig. 1a). To quantify the periodicity of Tpm3.1, we plotted fluorescence intensity profiles along regions in the AIS with visible periodicity and calculated the autocorrelation function for each profile. The average autocorrelation for the profiles measured showed an autocorrelation peak at a lag of 200 nm for both F-actin and Tpm3.1 (Fig. 1b and c, left panels). Owing to the pixel size of the camera used (40 nm), the distances recorded are multiples of 40. Accordingly, a lag of 200 nm corresponds to the ~190 nm reported earlier for actin rings and other components of the AIS sub-membranous lattice (D’Este et al., 2015; Leite et al., 2016; Leterrier et al., 2015; Xu et al., 2013; Zhong et al., 2014). For both F-actin and Tpm3.1, we measured the distance between individual neighboring fluorescence intensity peaks in each profile (Fig. 1b and c, right panels). The mean inter-peak distance for F-actin was 190.36 ± 1.7 nm (mean ± SEM), with 37.7% of the profiles 200 nm apart. The mean inter-peak distance for Tpm3.1 was 200.98 ± 1.5 nm (mean ± SEM), with 47.1% of the distances measuring 200 nm. In contrast, Tpm3.1 did not show clear periodicity in neither dendrites nor the distal regions of the axon (Supplementary Figure 1).

**Figure 1.**
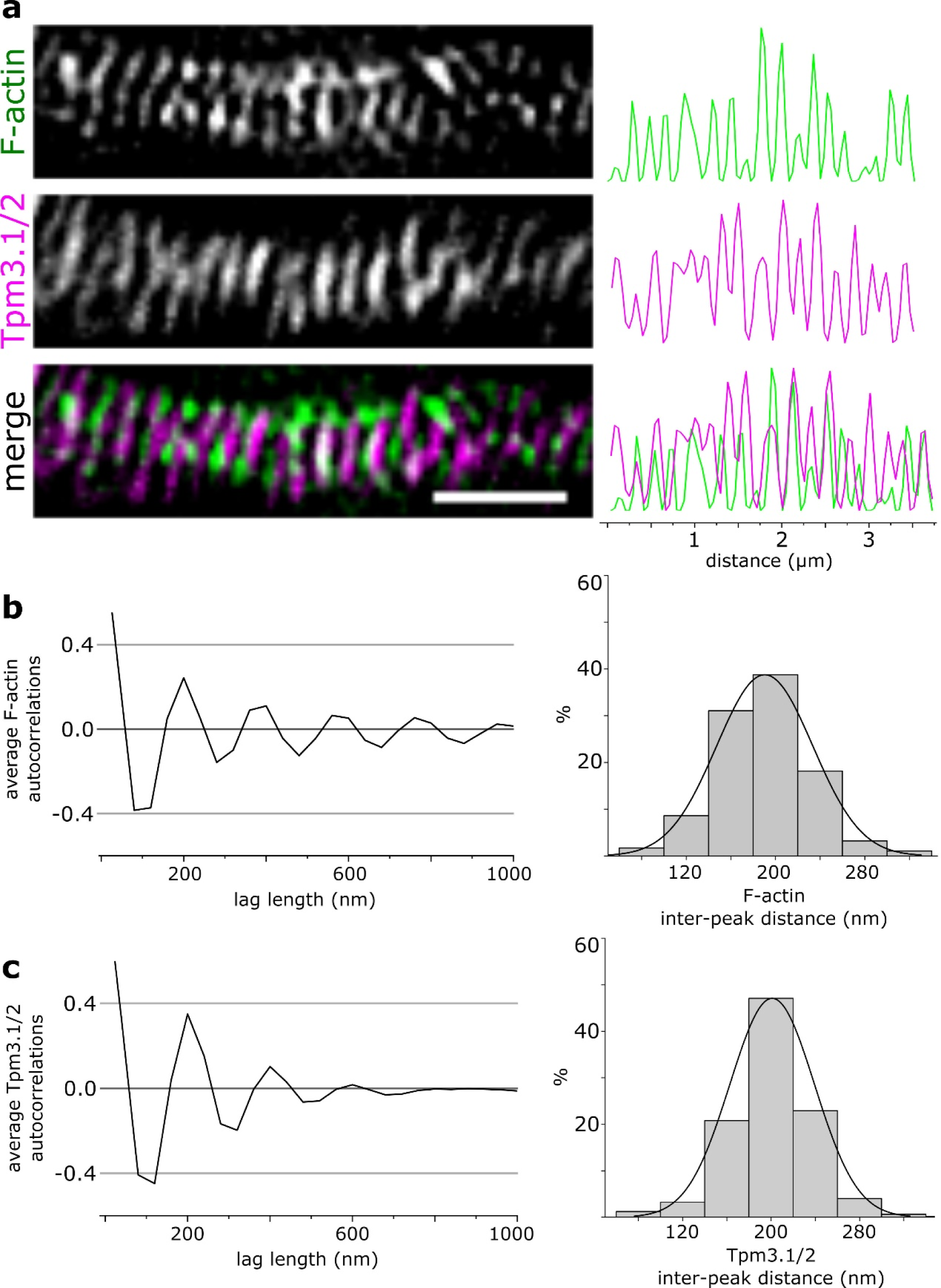
Tropomyosin isoform Tpm3.1 forms a periodic structure in the AIS. **a** SIM reconstruction of the AIS of a rat hippocampal neuron at 14 DIV labelled using anti-γ/9d and Alexa-488 tagged phalloidin to visualize Tpm3.1 and F-actin, respectively. Anti-Ankyrin G served to label the AIS. Tpm3.1/2 shows a periodic structure partially corresponding to actin rings in the AIS. Right: fluorescence intensity profile along the AIS. **b** Left: Average autocorrelation of normalized phalloidin fluorescence intensity profiles showing autocorrelation at 200 nm. Right: Distance between individual peaks in normalized phalloidin fluorescence intensity profiles. 37.7% of the peaks were separated by 200 nm. **c** Left: Average autocorrelation of normalized anti-γ/9d fluorescence intensity profiles showing autocorrelation at 200 nm. Right: Distance between individual peaks in normalized anti-γ/9d fluorescence intensity profiles. 47.1% of the peaks were separated by 200 nm. n = 25 cells, 4 independent experiments. Scale bar: 1 µm.

### The AIS contains patches of F-actin with a low rate of depolymerization

Previous studies revealed that, in addition to sub-membranous actin rings, actin forms patches in the AIS (Balasanyan et al., 2017; D’Este et al., 2015; Janssen et al., 2017; Jones et al., 2014; Leite et al., 2016; Leite and Sousa, 2016; Leterrier et al., 2015; Watanabe et al., 2012; Xu et al., 2013; Zhong et al., 2014) that may play a role in presynaptic terminals (D’Este et al., 2015) or in the filtering of vesicles carrying somatodendritic cargo (Balasanyan et al., 2017; Janssen et al., 2017; Watanabe et al., 2012). However, little is known about the dynamics and regulation of these actin patches. To investigate actin regulation, we expressed photoactivatable GFP-tagged actin (PAGFP-actin) in cultured rat hippocampal neurons to study the dynamics of F-actin in the AIS (Koskinen and Hotulainen, 2014; Patterson and Lippincott-Schwartz, 2002). We co-transfected neurons at 8-10 days *in vitro* (DIV) using mCherry and PAGFP-actin and imaged them 40-56 hours later. To label the AIS, we used an antibody against the extracellular domain of NF-186, 1-2 hours before imaging (Hedstrom et al., 2008). To visualize the distribution of F-actin in the AIS, we applied a brief 405-nm laser pulse within a 30 µm-long region along the AIS (Fig. 2a). The fluorescence intensity within this region was monitored for 3 minutes by capturing a frame every 3 seconds. Due to the fast rate of diffusion of free actin monomers, the first frame taken after photoactivation (0s) enables the visualization of only those monomers that were immobilized by incorporation into an actin filament (Honkura et al., 2008).

**Figure 2.**
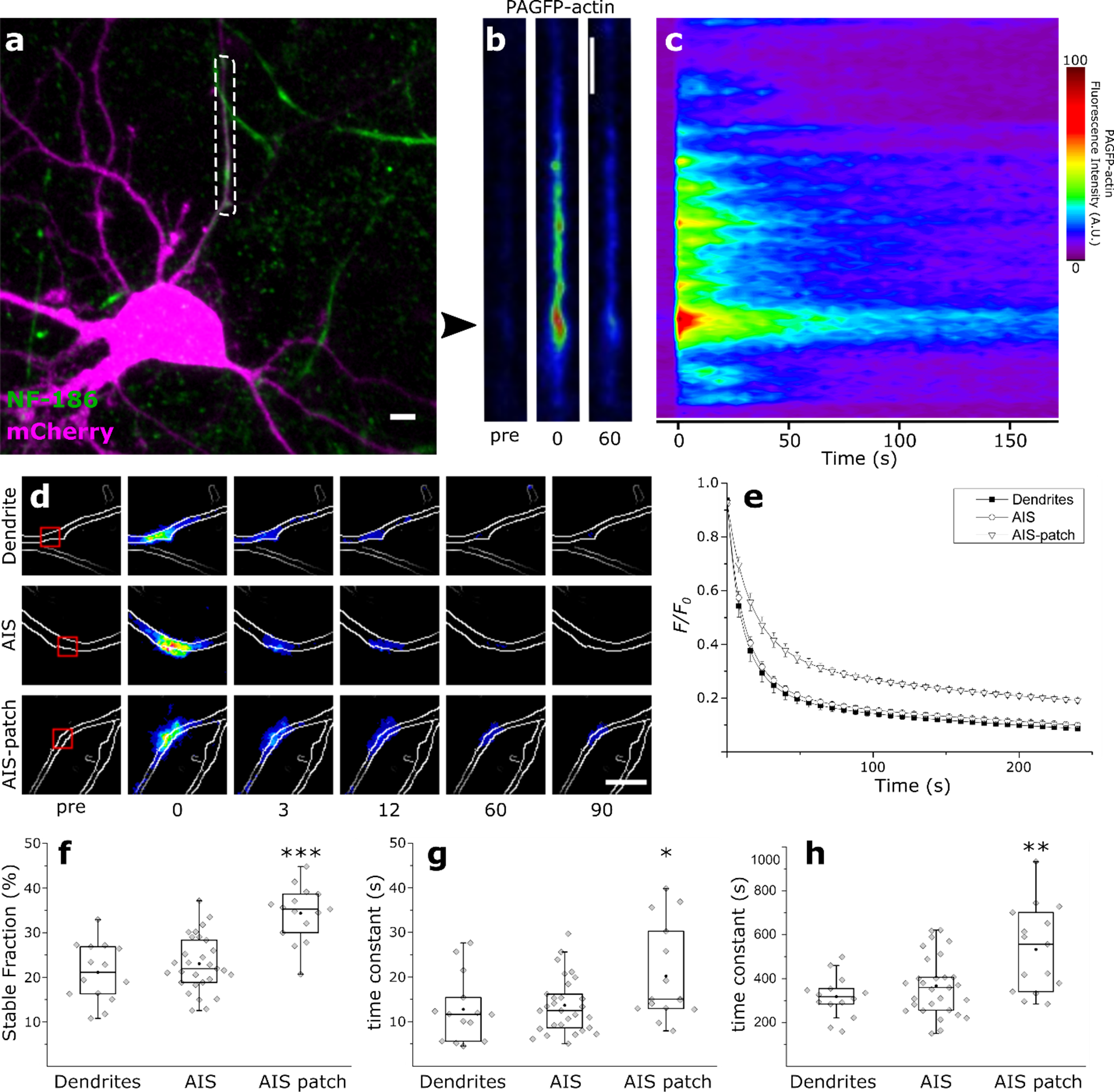
F-actin patches in the AIS have a lower rate of depolymerization. **a** We performed photoactivation within the dashed-box representing the entire AIS in rat hippocampal neurons expressing mCherry and PAGFP-actin and monitored PAGFP fluorescence over time. PanNF186 served to label the AIS. **b** Higher magnification of the dashed-box in (**a**) showing PAGFP-actin fluorescence 3 seconds before, immediately after, and 60 seconds after photoactivation. Arrowhead indicates F-actin patch. **c** PAGFP-actin fluorescence intensity profile along the AIS over time. **d** We performed photoactivation in a dendrite, the AIS, or an F-actin patch in the AIS (‘AIS patch’). Photoactivation was limited to the small boxed region to enable a more accurate measurement of F-actin dynamics. Contour lines were constructed using mCherry fluorescence. **e** Average normalized fluorescence decay curve fits over time in dendrites, the AIS, and F-actin patches in the AIS. We fit fluorescence decay curves to a double-exponential decay function and compared the fitting parameters across groups. **f** Percentage of the stable fraction in dendrites, the AIS, and AIS actin patches (ANOVA, Tukey’s test). **g** Time constants of the dynamic fractions (Mann-Whitney *U* test). **h** Time constants of the stable fractions (Mann-Whitney *U* test). Black circles represent mean value. Box borders represent the 25^th^ and 75^th^ percentiles, whiskers represent minimum and maximum values less than 1.5x the interquartile range lower or higher than the 25^th^ or 75^th^ percentiles, respectively (Tukey style). Dendrites: n = 14, 4 independent experiments; AIS: n = 29, 6 independent experiments; AIS patch: n = 15, 7 independent experiments. * denotes statistical significance. *: p < 0.05; **: p < 0.01; ***: p < 0.001. Scale bar: 5 µm.

The distribution of F-actin in the AIS was uneven and a prominent patch under 1 µm in diameter showed a higher fluorescence intensity, corresponding to a higher concentration of F-actin (Fig. 2b). Relative to the rest of the AIS, this actin patch was also the most long-lived (Fig. 2c). To measure the rate of depolymerization more accurately, we confined the photoactivation to a square area roughly 5 µm^2^ in size (Fig. 2d, red box). In addition to allowing for faster photoactivation, minimizing the area of photoactivation also minimizes the interference of photoactivated monomers that are incorporated into neighboring filaments after dissociation, leading to improved accuracy. Photoactivation was carried out within an AIS actin patch, in the AIS outside actin patches, and in a comparable dendritic segment that does not contain dendritic spines or branching points. An image was taken every 3 seconds and fluorescence intensity values were recorded. After subtracting the background fluorescence, we normalized the intensity values to the value at 0 s to obtain a normalized fluorescence decay curve. A double-exponential decay function gave the best fit for the decay curves in all groups (Koskinen and Hotulainen, 2014), indicating the presence of two pools of actin filaments with different rates of depolymerization. Accordingly, we fit the fluorescence decay curves to a double-exponential decay function (Fig. 2e) and the fitting parameters were compared across groups.

The average proportion of the stable fraction of actin filaments (Fig. 2f) was not significantly different between dendrites (21.1 ± 1.8%, mean ± SEM, n = 14, 4 independent experiments) and regions in the AIS outside the patches (23.0 ± 1.2%, mean ± SEM, n = 29, 6 independent experiments). Actin patches, however, had a higher proportion of stable filaments (34.4 ± 1.6%, mean ± SEM, n = 15, 7 independent experiments, p < 0.001, ANOVA, Tukey’s test). In contrast, using the same experimental setup we found the proportion of stable actin filaments in dendritic spines to be 18% and 30% in cultured hippocampal neurons at 14 and 21 DIV, respectively (Koskinen et al., 2014). Fig. 2g and h show the time constants for the dynamic and the stable pools of actin filaments, respectively. The average decay time constant of the dynamic pool was not significantly different between dendrites (12.76 ± 1.99 s, mean ± SEM, n = 14) and regions in the AIS outside the patches (13.66 ± 1.16 s, mean ± SEM, n = 29). The decay time constant of the dynamic pool was higher in the actin patches (20.2 ± 2.74 s, mean ± SEM, n = 15, p < 0.05, Mann-Whitney *U* test). The average decay time constant for the stable pool was not significantly different between dendrites (318 ± 25 s, mean ± SEM, n = 14) and regions in the AIS outside the patch (367 ± 25 s, mean ± SEM, n = 29). This value was higher in the actin patches (533 ± 50 s, mean ± SEM, n = 15, p < 0.01, Mann-Whitney *U* test) indicating that filaments in the AIS actin patches have a lower rate of depolymerization. It is likely that sub-membranous actin rings in the AIS had a minimal contribution to the readout in these experiments. This is partly due to the relatively low amount of F-actin in the sub-membranous lattice compared to intracellular F-actin within the area activated (10-fold less overall F-actin, Supplementary Figure 2). In addition, actin filaments in the sub-membranous rings are relatively stable (Abouelezz et al., 2019), possibly leading to a low rate of incorporation of PAGFP-actin monomers available for photoactivation. These data indicate that the AIS contains patches of F-actin that have a high proportion of stable filaments with a low rate of depolymerization.

Furthermore, we used PAGFP-actin to locate an actin patch with a low rate of depolymerization, then used anti-γ/9d to examine Tpm3.1/2 distribution. Tpm3.1/2 immunofluorescence showed a high intensity at the actin patch (Fig. 3b, arrowheads), indicating that Tpm3.1 colocalizes with actin patches in the AIS. Indeed, Tpm3.1/2 immunofluorescence in the AIS displays a patchy, non-uniform appearance resembling that of F-actin (Supplementary Figure 3). As the anti-γ/9d antibody simultaneously binds isoforms Tpm3.1 and the closely related (sharing 7 of 8 exons) isoform Tpm3.2, we separately briefly expressed YFP-tagged Tpm3.1 and Tpm3.2 in cultured rat hippocampal neurons and examined their distribution. Anti-ankyrin G served to label the AIS. YFP-Tpm3.2 showed a faint and diffuse distribution in the AIS and distal axon, and mild enrichment in dendritic spines (Supplementary Figure 4b). Conversely, we observed bright puncta of YFP-Tpm3.1 in the AIS and the distal axon, and a diffuse distribution in the somatodendritic domain (Supplementary Figure 4a). These puncta had a similar distribution to AIS actin patches and were resistant to detergent extraction (Supplementary Figure 5), indicating cytoskeletal association.

**Figure 3.**
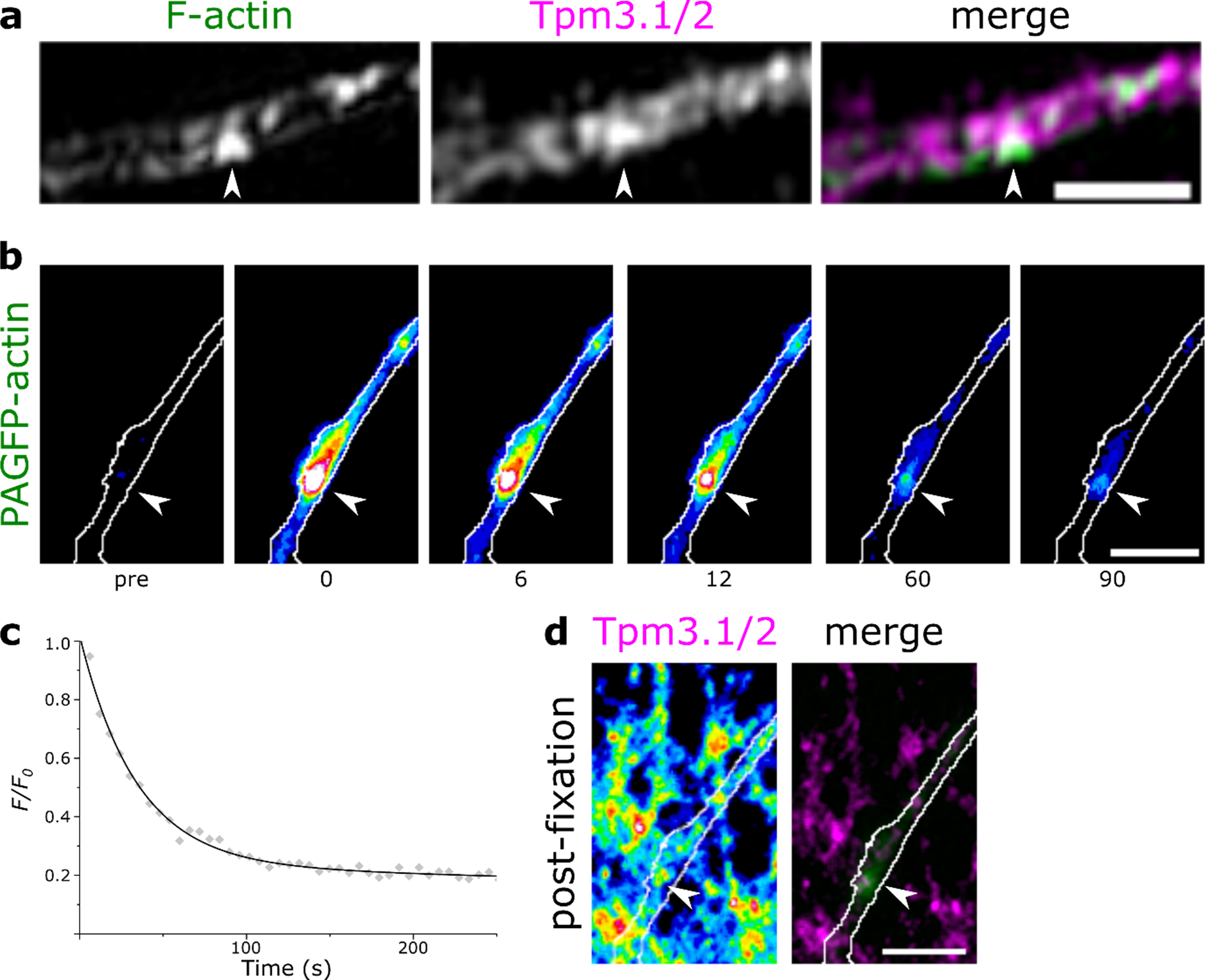
Tropomyosin Tpm3.1 decorates actin patches in the AIS. **a** Maximum intensity projection of 3D-SIM reconstructions for F-actin and Tpm3.1/2. Arrowheads indicate actin patch. Scale bar: 1 µm. **b** Actin patch visualized in live hippocampal neuron using PAGFP-actin before photoactivation (pre), immediately after activation (0 s) and the time-points indicated in seconds. Arrowheads indicate actin patch. **c** Fluorescence decay over time (gray diamonds) of the actin patch in (**b**) and a double-exponential decay fit (solid black line). **d** Tpm3.1/2 distribution visualized using anti-γ/9d in the same area after fixation in 4% PFA. The intensity of Tpm3.1/2 immunofluorescence was higher in the region corresponding to the actin patch visualized in (**b**) (arrowhead). Scale bar: 5 µm.

Based on these data, we conclude that Tpm3.1 is present in the AIS and colocalizes with actin patches. Importantly, Tpm3.1 is not exclusively localized to the AIS. As earlier studies show that Tpm3.1 increases the F-actin pool and lowers the rate of actin depolymerization (Broschat, 1990; Schevzov et al., 2008), it is plausible that Tpm3.1 similarly increases the concentration of F-actin and lowers the rate of actin depolymerization in AIS actin patches.

### Tpm3.1 is required for maintaining the structure of the AIS

We next examined the consequences of the perturbation of Tpm3.1 function for AIS structure. We used two distinct, well-characterized, small-molecule Tpm3.1 inhibitors, namely TR100 (Bonello et al., 2016; Kee et al., 2018; Kee et al., 2015; Stehn et al., 2013) and Anisina (ATM3507) (Currier et al., 2017; Stehn et al., 2016) to inhibit Tpm3.1 and examined the accumulation of ankyrin G and other AIS markers at the AIS in mature cultured rat hippocampal neurons. TR100-mediated inhibition of Tpm3.1 does not inhibit the binding of Tpm3.1 to actin filaments, but negates its effect on the rate of depolymerization (Bonello et al., 2016). The effects of TR100 on glucose-stimulated insulin secretion were not seen in Tpm3.1 knockout mice, indicating its specificity (Kee et al., 2018).

We incubated sparse cultures of rat hippocampal neurons at 10 DIV using DMSO (0.2%), TR100 (10 or 15 µM), or Anisina (5 or 7.5 µM) for 2, 3, or 6 hours and then used anti-ankyrin G to visualize the distribution of ankyrin G. Anti-MAP2 served to label the somatodendritic domain. Both TR100- and Anisina-mediated Tpm3.1 inhibition led to a notable reduction in the accumulation of ankyrin G at the AIS (Fig. 4a). To quantify the reduction in ankyrin G accumulation, we blindly traced the fluorescence intensity profile of ankyrin G along the initial 30 µm of every neurite to calculate an AIS localization index (ALI, see methods) (Fig. 4b). For all treatment conditions, the ALI was lower than the corresponding DMSO controls (Mann-Whitney *U* test; DMSO 0.2%, 2 hours: 0.94 ±0.006, mean ± SEM; DMSO 0.2%, 3 hours: 0.93 ± 0.006; DMSO 0.2%, 6 hours: 0.96 ± 0.005; TR100 10 µM, 2 hours: 0.59 ± 0.050, p < 0.001; TR100 10 µM, 3 hours: 0.50 ± 0.051, p < 0.001; TR100 10 µM, 6 hours: 0.42 ± 0.066, p < 0.001; TR100 15 µM, 2 hours: 0.45 ± 0.060, p < 0.001; TR100 15 µM, 3 hours: 0.42 ± 0.060, p < 0.001; TR100 15 µM, 6 hours: 0.35 ± 0.056, p < 0.001; Anisina 5 µM, 2 hours: 0.58 ± 0.055, p < 0.001; Anisina 5 µM, 3 hours: 0.49 ± 0.049, p < 0.001; Anisina 5 µM, 6 hours: 0.44 ± 0.039, p < 0.001; Anisina 7.5 µM, 2 hours: 0.51 ± 0.041, p < 0.001; Anisina 7.5 µM, 3 hours: 0.41 ± 0.062, p < 0.001; Anisina 7.5 µM, 6 hours: 0.36 ± 0.075, p < 0.001; for each treatment, n = 12, 3 independent experiments) (Fig. 4c). The mean ALI was negatively correlated with the concentration of both TR100 (Pearson’s coefficient; 2 hours: −1.0; 3 hours: −0.98; 6 hours: −0.97) and Anisina (Pearson’s coefficient; 2 hours: - 0.99; 3 hours: −0.98; 6 hours: −0.98), as well as with the duration of the treatment (Pearson’s coefficient; TR100 10 µM: −0.94; TR100 15 µM: −1.0; Anisina 5 µM: −0.90; Anisina 7.5 µM: −0.90), suggesting dose- and time-dependence. We similarly observed a reduction in the mean ALI of ankyrin G after overnight inhibition of Tpm3.1 (Supplementary Figure 6). Tpm3.1 inhibition also abolished the accumulation at the AIS of all other AIS markers tested, namely TRIM46, EB1, and neurofascin-186 (Supplementary Figure 7).

**Figure 4.**
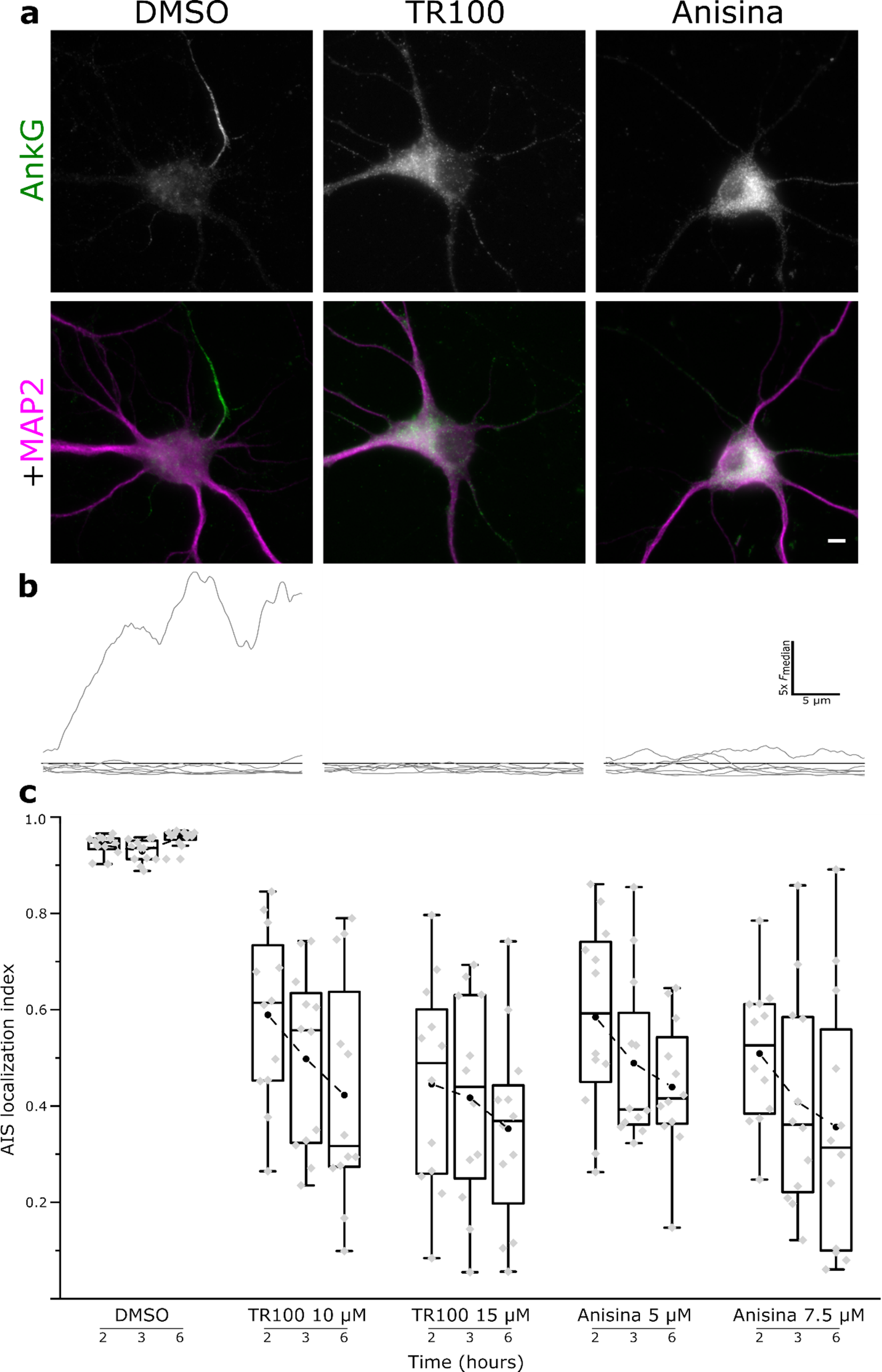
Inhibition of Tpm3.1 reduces the accumulation of ankyrin G at the AIS. **a** Rat hippocampal neurons treated at 10 DIV using DMSO or the small-molecule Tpm3.1 inhibitors TR100 or Anisina (ATM3507) for 2, 3, or 6 hours. Anti-MAP2 served to label the somatodendritic domain, anti-ankyrin G served to measure the accumulation of ankyrin G. **b** Smoothed ankyrin G fluorescence intensity line profiles (gray lines) along each neurite of the corresponding cell in (**a**), normalized to the median peak value (black line). **c** AIS localization indices for neurons treated using DMSO, TR100 (10 or 15 µM), or Anisina (5 or 7.5 µM) for 2, 3, or 6 hours. All treatment groups were significantly different from DMSO controls (Mann-Whitney *U* test). The mean ALI of the treatment groups was negatively correlated with treatment duration and concentration. Black circles represent mean values. Box borders represent the 25^th^ and 75^th^ percentiles, whiskers represent minimum and maximum values less than 1.5x the interquartile range lower or higher than the 25^th^ or 75^th^ percentiles, respectively (Tukey style). For each treatment: n = 12, 3 independent experiments. Dotted lines connect mean values. Scale bar: 5 µm.

While the loss of AIS structure upon Tpm3.1 inhibition using either TR100 or Anisina was clear, it is possible that this was a secondary effect rising from a general decay in neuronal health rather than a direct effect of the perturbation of Tpm3.1 function. In fact, oxidative stress (Clark et al., 2017) and neuronal injury (Schafer et al., 2009) resulted in the disruption of the AIS in a calpain-mediated manner. Schafer et al. (2009) showed that neuronal injury induced irreversible AIS disassembly through calpain-mediated proteolysis of ankyrin G and βIV-spectrin. However, this process was blocked by using the calpain inhibitor MDL28170. Thus, to test whether a similar calpain-dependent mechanism contributes to the loss of the AIS upon Tpm3.1 inhibition, we added calpain inhibitors to TR100 or Anisina treatments and measured the ALI of ankyrin G. Indeed, the loss of AIS structure upon Tpm3.1 inhibition was calpain-independent; the mean ALI for all treatment conditions did not significantly change in the presence of 100 µM of the calpain inhibitor MDL28170 for the entire duration of the treatment (Supplementary Figure 8).

To verify that the loss of ankyrin G accumulation at the AIS is a result of the loss of Tpm3.1 function, we generated a conditional knockout mouse model (Tp9 line, Supplementary Figure 9) for all Tpm3 isoforms containing exon 1b (namely Tpm3.1, Tpm3.2, Tpm3.3, Tpm3.4, Tpm3.5, Tpm3.7, Tpm3.8, Tpm3.9, Tpm3.12, and Tpm3.13). We plated dissociated hippocampal neurons from these mice onto PDL-coated coverslips. To induce protein depletion, we transduced cultures using CMV-EGFP-Cre adeno-associated viruses (UNC Vector Core Facility). This resulted in a 10% transduction efficiency leading to a mixture of wild-type and Tpm3-depleted neurons. Alternatively, we used CMV-EGFP adeno-associated viruses to transduce GFP— but not Cre—expression in sister cultures as controls. We confirmed the depletion of Tpm3 isoforms through immunostaining using an anti-Tpm3 antibody (Fig. 5a). We then used anti-ankyrin G to visualize the accumulation of ankyrin G at the AIS (Fig. 5b). The average relative fluorescence intensity of ankyrin G at the AIS in Tpm3-depleted neurons was lower (0.74 ± 0.04, mean ± SEM, n = 61, 3 independent experiments, p < 0.001, Mann Whitney *U* test) than in control neurons (1 ± 0.04, mean ± SEM, n = 63, 3 independent experiments) (Fig 5c). The effect of knocking out *Tpm3* was milder than what we expected. However, conventional *Tpm4* knockout mice (Tp16 line) (Pleines et al., 2017) did not show any detectable difference in the accumulation of ankyrin G at the AIS (Supplementary Figure 10). We speculate that other proteins— possibly other tropomyosin isoforms— compensated for the loss of Tpm3.1 function, leading to an effect milder than acute small-molecule inhibition.

**Figure 5.**
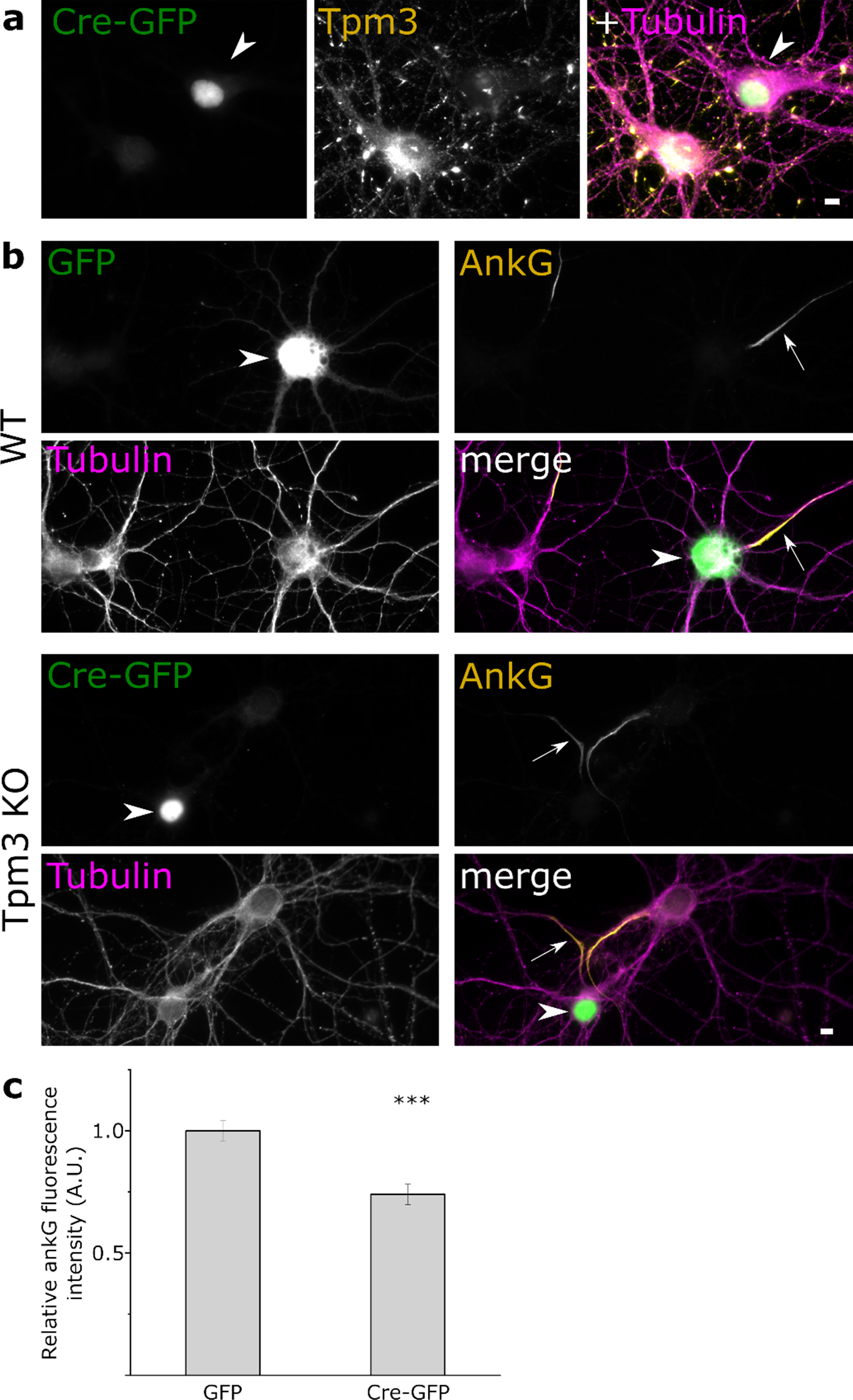
*Tpm3* conditional knockout neurons show a reduced accumulation of ankyrin G at the AIS. **a** Tpm3 immunofluorescence in cultured hippocampal neurons of *Tpm3* knockout mice. Arrowhead indicates a neuron expressing GFP-tagged Cre-recombinase (Cre-GFP). **b** Ankyrin G immunofluorescence for GFP-(wild-type, WT) and Cre GFP-expressing (*Tpm3* KO) neurons. β3-Tubulin was used to label neurons. Arrowheads indicate transfected neurons, arrows indicate axons of transfected neuron. **c** Average relative ankyrin G fluorescence intensity for each group (Mann-Whitney *U* test). Columns represent mean values. Error bars represent standard error of mean. GFP: n = 63, 3 independent experiments; Cre-GFP: n = 61, 3 independent experiments. * denotes statistical significance. ***: p < 0.001. Scale bar: 5 µm.

Furthermore, we obtained similar results by expressing shRNA specific to exon 9d for 4 days, which only depletes *Tpm3* isoforms Tpm3.1 and Tpm3.2 (Supplementary figure 11). Importantly, the effect was rescued by over-expressing YFP-tagged human Tpm3.1, which differs in sequence within the shRNA-targeting region (Supplementary figure 12). Thus, we conclude that both pharmacological inhibition and genetic depletion of Tpm3.1 cause a notable defect in AIS structure. Together, these data suggest that the reduced accumulation of ankyrin G at the AIS is the result of the loss of Tpm3.1 function, indicating that Tpm3.1 is necessary for maintaining the structure of the AIS.

### Tpm3.1 is necessary for maintaining the selectivity of axonal transport and sodium channel clustering at the AIS

Vesicular filtering and the diffusion barrier at the AIS require an intact actin cytoskeleton (Song et al., 2009; Winckler et al., 1999). Somatodendritic cargo entering the AIS halt at regions of high F-actin concentration, in a process that is dependent on myosin motors (Balasanyan et al., 2017; Janssen et al., 2017; Watanabe et al., 2012). To test if Tpm3.1 is required for maintaining the AIS vesicular filter, we fixed sparse cultures of rat hippocampal neurons at 10 DIV after an overnight treatment using DMSO (0.2%), LatB (5 µM), or TR100 (5 µM) and used antibodies against the somatodendritic glutamate receptor subunit GluA1 to visualize its distribution. Anti-MAP2 served to label the somatodendritic domain. Contrary to the somatodendritic localization of GluA1 observed in DMSO-treated neurons, we detected GluA1 immunofluorescence in both the dendrites and axons of treated neurons (Fig. 6a). We used maximum intensity projection images from confocal stacks to blindly measure the mean fluorescence intensity along the axon and dendrites to calculate the axon-to-dendrite ratio for each group (Lewis et al., 2009). LatB-treated neurons showed a higher axon-to-dendrite ratio (0.38 ± 0.02, mean ± SEM, n = 15 neurons, 2 independent experiments, p < 0.01, Mann-Whitney *U* test) than DMSO-treated neurons (0.25 ± 0.03, mean ± SEM, n = 14 neurons, 2 independent experiments). TR100-treated neurons also showed a higher axon-to-dendrite ratio (0.49 ± 0.05, mean ± SEM, n = 18 neurons, 2 independent experiments, p < 0.001, Mann-Whitney *U* test) (Fig. 6b). This suggests that Tpm3.1 function is necessary for maintaining the selectivity of axonal transport at the AIS. In addition, we wanted to examine the effect of Tpm3.1 inhibition on the clustering of sodium channels, which is essential for spike generation at the AIS (Kole et al., 2008). This accumulation of channels is achieved through interactions with ankyrin G (Jenkins and Bennett, 2001; Zhou et al., 1998). We used antibodies against voltage-gated sodium channels (panNa_v_) and the somatodendritic marker MAP2 to visualize sodium channel clustering at the AIS in sparse cultures of rat hippocampal neurons at 10 DIV after an overnight treatment using DMSO (0.2%), LatB (5 µM), or TR100 (5 µM). DMSO- and LatB-treated neurons showed clear detectable clustering of panNa_v_ immunofluorescence in the AIS, while TR100-treated neurons displayed a relatively homogenous distribution in all neurites (Fig. 7a). To quantitatively examine this effect, we used maximum intensity projections of confocal stacks to blindly record panNa_v_ immunofluorescence along the initial 30 µm of each neurite to calculate the ALI (Fig. 7b). There was no difference in the mean ALI between DMSO-treated neurons (0.76 ± 0.02, mean ± SEM, n = 17 neurons, 3 independent experiments) and LatB-treated neurons (0.75 ± 0.03, mean ± SEM, n = 17 neurons, 3 independent experiments). In contrast, TR100-treated neurons showed a lower ALI (0.37 ± 0.04, mean ± SEM, n = 17 neurons, 3 independent experiments, p < 0.001, Mann-Whitney *U* Test), indicating a more homogenous distribution across neurites (Fig. 7c). These data suggest that Tpm3.1 is required for the clustering of sodium channels at the AIS.

**Figure 6.**
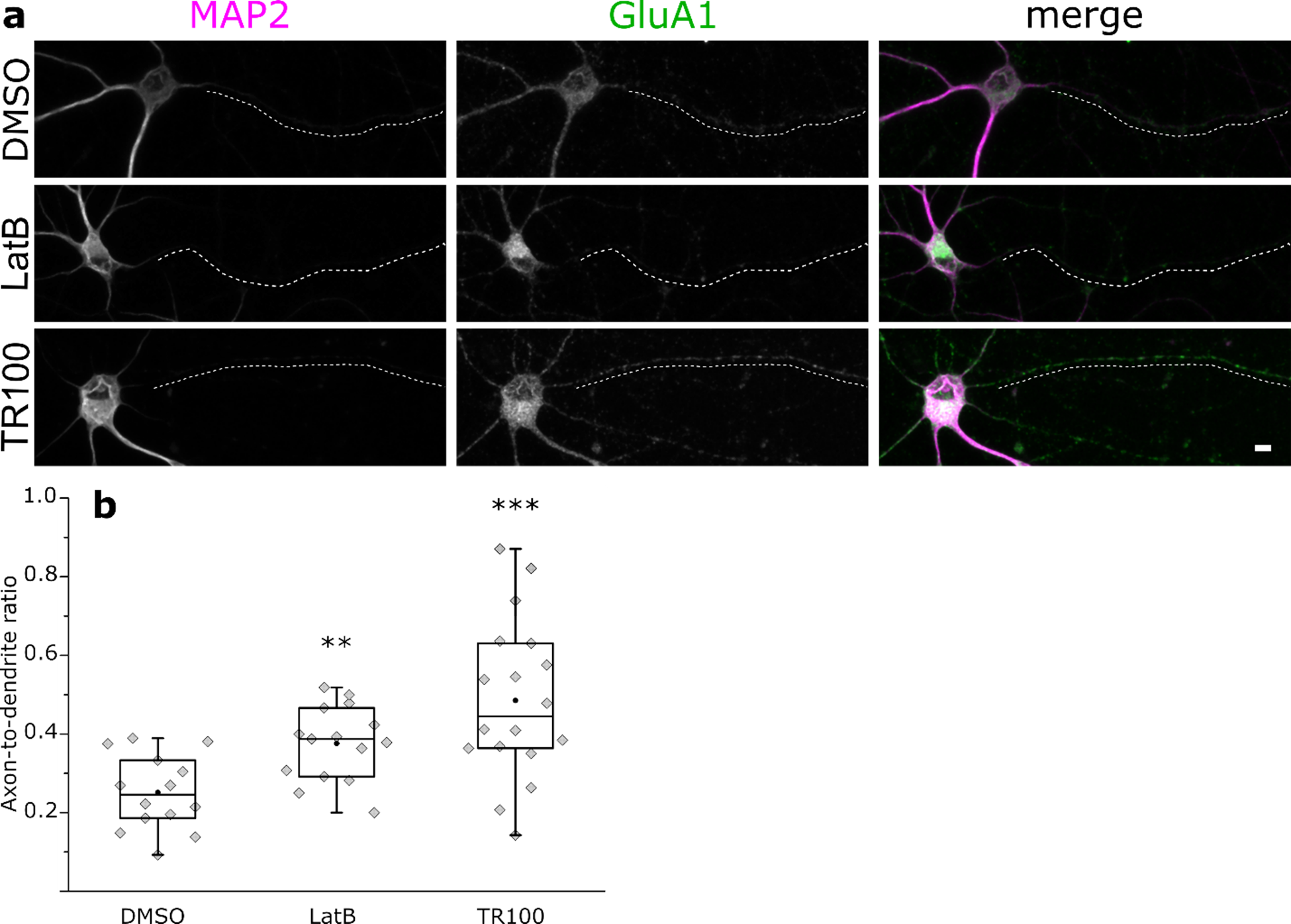
Tpm3.1 inhibition leads to the redistribution of the somatodendritic marker GluA1. **a** MAP2 and GluA1 immunofluorescence in rat hippocampal neurons incubated overnight at 9-11 DIV in DMSO, LatB, or TR100. Dashed lines represent axons. **b** GluA1 axon-to-dendrite ratios were higher in LatB- and TR100-treated neurons (Mann-Whitney *U* test). Black circles represent mean value. Box borders represent the 25^th^ and 75^th^ percentiles, whiskers represent minimum and maximum values less than 1.5x the interquartile range lower or higher than the 25^th^ or 75^th^ percentiles, respectively (Tukey style). DMSO 0.2%: n = 14, 2 independent experiments; LatB 5 µM: n = 14, 2 independent experiments; TR100 5 µM: n = 18, 2 independent experiments. * denotes statistical significance. **: p < 0.01; ***: p < 0.001. Scale bar: 5 µm.

**Figure 7.**
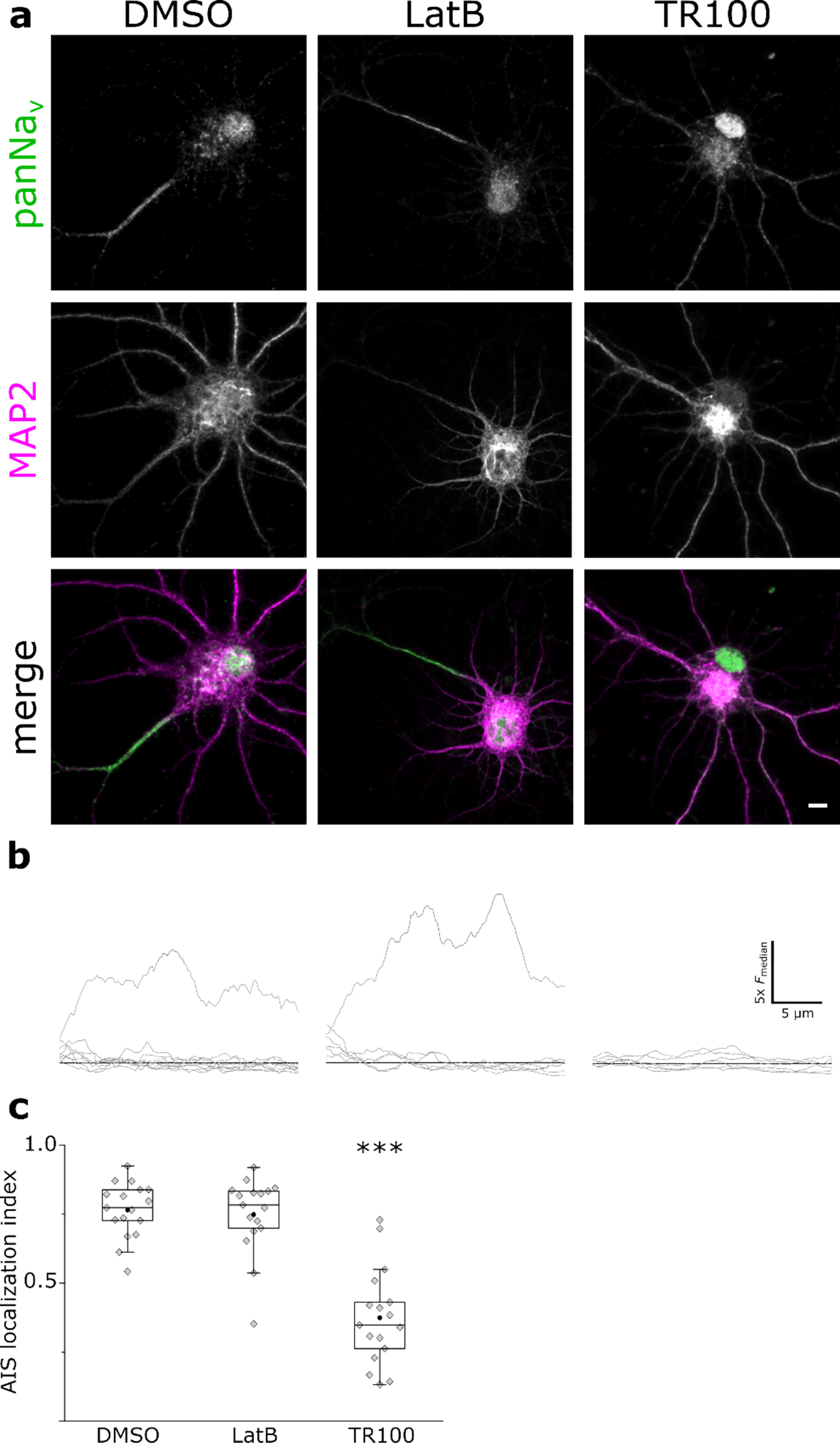
Tpm3.1 inhibition leads to the loss of voltage-gated sodium channels clustering at the AIS. **a** MAP2 and panNa_v_ immunofluorescence in rat hippocampal neurons incubated overnight at 9-11 DIV in DMSO, LatB, or TR100. **b** Smoothed panNa_v_ fluorescence intensity line profiles (gray lines) along each neurite of the corresponding neuron in (**a**) normalized to the median value (black line). **c** AIS localization indices for each group (Mann-Whitney *U* Test). Black circles represent mean value. Box borders represent the 25^th^ and 75^th^ percentiles, whiskers represent minimum and maximum values less than 1.5x the interquartile range lower or higher than the 25^th^ or 75^th^ percentiles, respectively (Tukey style). DMSO 0.2%: n = 17, 3 independent experiments; LatB 5 µM: n = 17, 3 independent experiments; TR100 5 µM: n = 17, 3 independent experiments. * denotes statistical significance. ***: p < 0.001. Scale bar: 5 µm.

### Tpm3.1 inhibition leads to a reduction in firing frequency

The initiation of action potentials is facilitated at the AIS by the clustering of ion channels (Kole et al., 2008), and is dependent on an intact AIS structure (Leterrier et al., 2017). To examine the effect of Tpm3.1 inhibition on spike generation, we recorded the activity of cultured rat hippocampal neurons at 16-18 DIV in current-clamp experiments in the presence of either DMSO (0.2%) or Anisina (2.5 µM). We introduced depolarizing steps of 100-200 pA at 10 s intervals and monitored the firing frequency 2 and 15 minutes after the introduction of DMSO or Anisina (Fig. 8a). The mean firing frequency of DMSO-treated neurons remained unchanged 15 minutes after the introduction of DMSO (at 2 minutes: 18.3 Hz ± 2.9, mean ± SEM; at 15 minutes: 20.9 Hz ± 4.2, mean ± SEM, n = 7 neurons, 5 independent experiments, paired-sample *t*-test). Conversely, Anisina-treated neurons showed a significant attenuation of firing frequency after 15 minutes (at 2 minutes: 21.7 Hz ± 4.2, mean ± SEM; at 15 minutes: 16.7 Hz ± 2.6, mean ± SEM, n = 7 neurons, 5 independent experiments, p < 0.05, paired-sample *t*-test). The change in mean firing frequency 15 minutes after introducing DMSO (9.5% ± 7.0, mean ± SEM) was significantly different from that of neurons treated using Anisina (−20.9% ± 3.4, mean ± SEM, p < 0.01, two-sample *t*-test, Fig. 8b). These data indicate that Tpm3.1 is required for maintaining AIS function in the initiation of action potentials, consistent with the loss of sodium channel clustering at the AIS.

**Figure 8.**
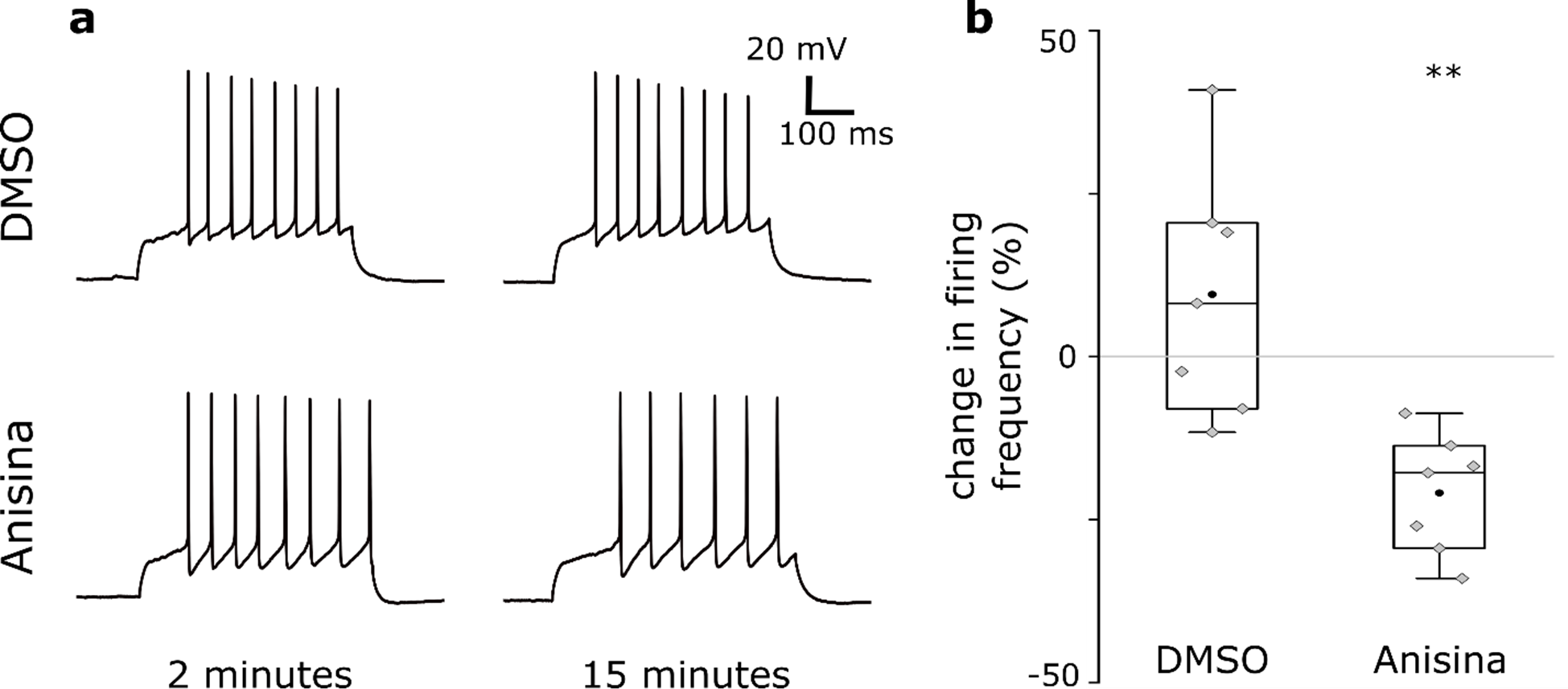
Tpm3.1 inhibition leads to a reduction in firing frequency. **a** Individual traces from current-clamp (depolarizing step of 100 pA for 500 ms) recordings of rat hippocampal neurons in culture 2 and 15 minutes after treatments using either DMSO (0.2%) or Anisina (2.5 µM). **b** The percentage change in firing frequency 15 minutes after introducing each treatment relative to the firing frequency at 2 minutes. Anisina-mediated inhibition of Tpm3.1 led to the attenuation of firing frequency 15 minutes after introduction. Black circles represent mean value. Box borders represent the 25^th^ and 75^th^ percentiles, whiskers represent minimum and maximum values less than 1.5x the interquartile range lower or higher than the 25^th^ or 75^th^ percentiles, respectively (Tukey style). DMSO 0.2%: n = 7, 5 independent experiments; Anisina 2.5 µM: n = 7, 5 independent experiments. * denotes statistical significance. **: p < 0.01.

### Tpm3.1 inhibition leads to the gradual collapse of the AIS actin cytoskeleton

Our results show that Tpm3.1 is important for the accumulation of ankyrin G at the AIS, but it is not clear how Tpm3.1 inhibition leads to the loss of ankyrin G accumulation. We are not aware of any reports suggesting direct interaction between Tpm3.1 and ankyrin G. Thus, we hypothesized that the loss of Tpm3.1 function adversely affects the overall structure and organization of the actin cytoskeleton in the AIS. This disorganization would then ultimately lead to the loss of ankyrin G accumulation, which would be sufficient for the loss of other AIS markers (Hedstrom et al., 2008; Jenkins and Bennett, 2001; Zhou et al., 1998).

To test whether Tpm3.1 inhibition disrupts actin structures in the AIS, we employed super-resolution microscopy techniques to examine the periodicity of F-actin in the AIS. We treated sparse cultures of rat hippocampal neurons at 14 DIV using DMSO (0.2%), TR100 (10 µM), or Anisina (5 µM) for 6 hours. In addition, we treated cultures at DIV 13 using LatB (5 µM) overnight, similar to Winckler et al. (1999). Consistent with our earlier report (Abouelezz et al., 2019), LatB-treated neurons showed an overall lower phalloidin fluorescence intensity, reflecting a decrease in overall F-actin (data not shown). Periodic actin rings were visible for all groups, indicating the persistence of the sub-membranous lattice, even in the absence of ankyrin G (Fig. 9a). We blindly plotted fluorescence intensity profiles in regions within the AIS where periodicity was visible (DMSO: n = 13 neurons, 4 independent experiments; LatB: n = 13 neurons, 4 independent experiments; TR100: n = 13 neurons, 3 independent experiments; Anisina: n = 13 neurons, 3 independent experiments) and calculated the autocorrelation function. All groups showed autocorrelation at a lag of 200 nm (Fig. 9b). Owing to the pixel size of the camera used (40 nm), the distances recorded are multiples of 40. Accordingly, a lag of 200 nm corresponds to the ~190 nm reported earlier for actin rings and other components of the AIS sub-membranous lattice (D’Este et al., 2015; Leite et al., 2016; Leterrier et al., 2015; Xu et al., 2013; Zhong et al., 2014). In addition, we blindly measured the distance between individual peaks in each fluorescence intensity profile and compared the distribution of the inter-peak distances across groups (Fig 9c). 54% of the inter-peak distances in DMSO-treated neurons were 200 nm, while the mean inter-peak distance was 192.49 ± 1.37 nm, mean ± SEM. The distribution of the inter-peak distances in LatB-treated neurons was not significantly different from DMSO controls, with 51.2% of the inter-peak distances at 200 nm, and a mean inter-peak distance of 188.7 ± 1.38 nm, mean ± SEM (p = 0.47, Kolmogorov Smirnov test). In contrast, TR100-treated neurons showed a less uniform distribution with only 41.7% of the inter-peak distance at 200 nm, and a mean inter-peak distance of 182.7 ± 1.74 nm, mean ± SEM (p < 0.01, Kolmogorov Smirnov test). Similarly, only 39.8% of the distances measured in Anisina-treated neurons were 200 nm, with a mean inter-peak distance of 189.0 ± 1.94 nm, mean ± SEM, a distribution significantly different from DMSO controls (p < 0.01, Kolmogorov Smirnov test). We also obtained similar results using stochastic optical reconstruction microscopy (STORM) (Supplementary figure 13).

**Figure 9.**
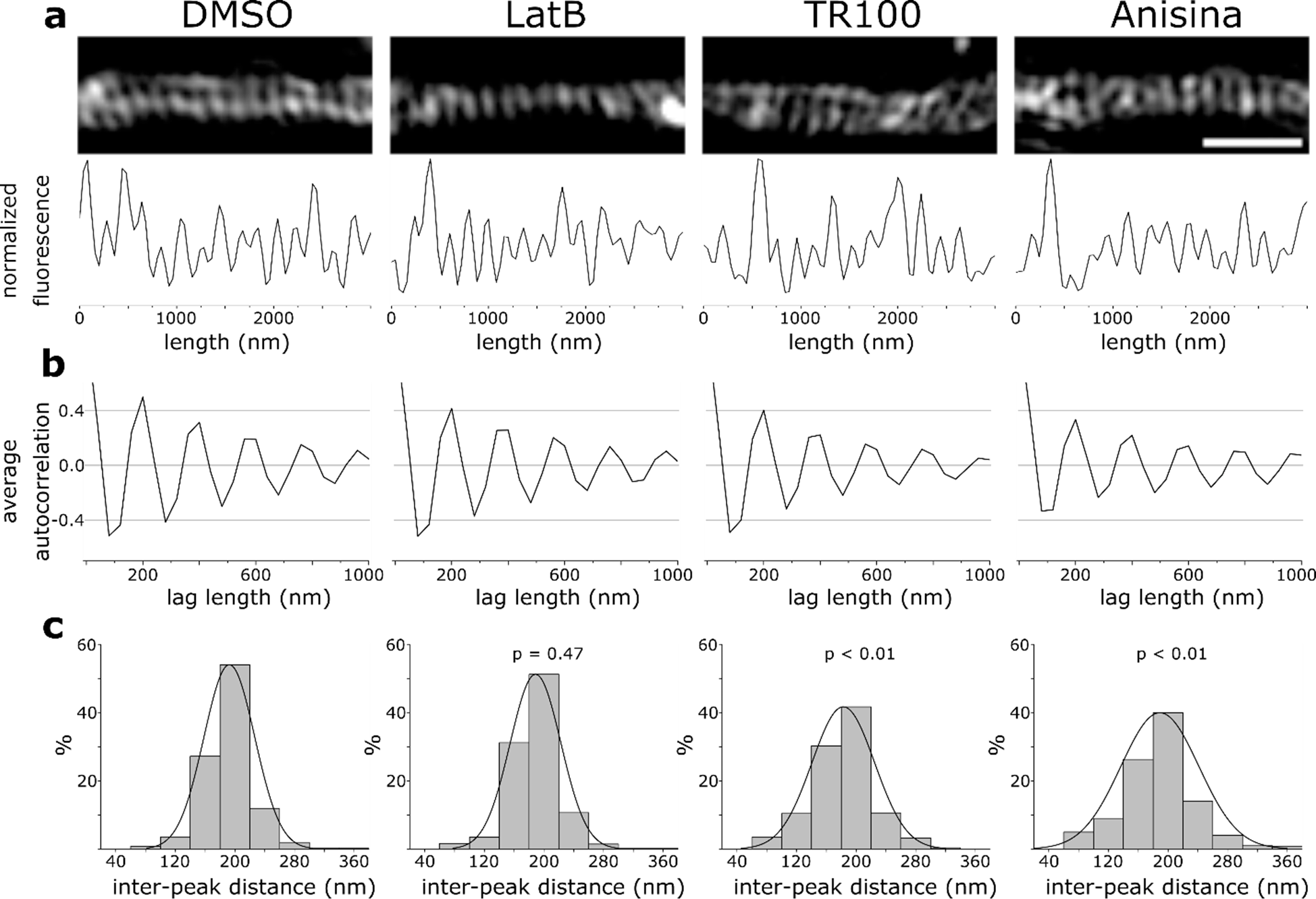
Tpm3.1 inhibition disrupts the periodicity of actin rings in the AIS. **a** SIM reconstructions of F-actin in the AIS of neurons treated at 14 DIV using DMSO, LatB, TR100, or Anisina (ATM3507), visualized using Alexa-488 tagged phalloidin. **b** Average autocorrelation of normalized fluorescence intensity profiles showing autocorrelation at 200 nm for all groups. **c** Distribution of distances between individual peaks in fluorescence intensity profiles for each group. The distribution of inter-peak distances in TR100- and Anisina-treated neurons was significantly different (p < 0.01) from DMSO- and LatB-treated neurons (Kolmogorov-Smirnov test). DMSO: n = 13 neurons, 4 independent experiments; LatB: n = 13 neurons, 4 independent experiments; TR100: n = 13 neurons, 3 independent experiments; Anisina: n = 13 neurons, 3 independent experiments. p values are relative to DMSO (Kolmogorov Smirnov test). Scale bar: 1 µm.

In conclusion, we found that the periodicity of sub-membranous actin rings in the AIS was resistant to LatB. As LatB disrupts the actin cytoskeleton through sequestering free actin monomers (thus inhibiting actin polymerization), stable actin filaments with a low rate of depolymerization may be less susceptible to LatB. In contrast, the inhibition of Tpm3.1 for the duration of the experiments (6 hours) disrupted—but did not entirely abolish—the periodicity of sub-membranous actin rings. In addition to changing the uniformity of the periodicity, visual inspection revealed that actin rings were often tilted after Tpm3.1 inhibition, losing their nature of parallel transverse stripes (Fig. 9a). In line with this, Tpm3.1 inhibition led to a reduction in the frequency of actin patches in the AIS compared to DMSO-treated neurons (DMSO: 0.58 ± 0.06 patches/µm, mean ± SEM, n = 13 neurons, 3 independent experiments; LatB: 0.71 ± 0.04 patches/µm, mean ± SEM, n = 13 neurons, 3 independent experiments, p = 0.19; TR100: 0.42 ± 0.02 patches/µm, mean ± SEM, n = 13 neurons, 3 independent experiments, p < 0.05; Anisina: 0.3 ± 0.05 patches/µm, mean ± SEM, n = 12 neurons, 3 independent experiments, p < 0.001, ANOVA, Tukey’s test).

Finally, we examined the effect of perturbing Tpm3.1 in cultured neurons on myosin IIB. Tpm3.1 recruits and activates myosin IIB (Bryce et al., 2003; Gateva et al., 2017), and recent work has revealed an important role for myosin II in AIS structure (Berger et al., 2018; Evans et al., 2017; Wang et al., 2018). We expressed Cre-GFP in cultured hippocampal neurons of *Tpm3* conditional knockout mice (Tp9 line) using either viral transduction or lipofection and used anti-myosin IIB to examine myosin IIB distribution (Fig. 10). Neurons expressing Cre-GFP after viral transduction showed a lower intensity of myosin IIB immunofluorescence (0.72 ± 0.1, mean ± SEM, n = 12) relative to neighboring control neurons (1 ± 0.06, mean ± SEM, p < 0.05, two-sample *t*-test, n = 22, 3 independent experiments). Similarly, compared to neighboring control neurons (1 ± 0.05, mean ± SEM, n = 31), neurons expressing Cre-GFP after lipofection showed a lower intensity of myosin IIB immunofluorescence (0.81 ± 0.06, mean ± SEM, p < 0.05, two-sample *t*-test, n = 12, 3 independent experiments). The results are summarized in Fig. 10.

**Figure 10.**
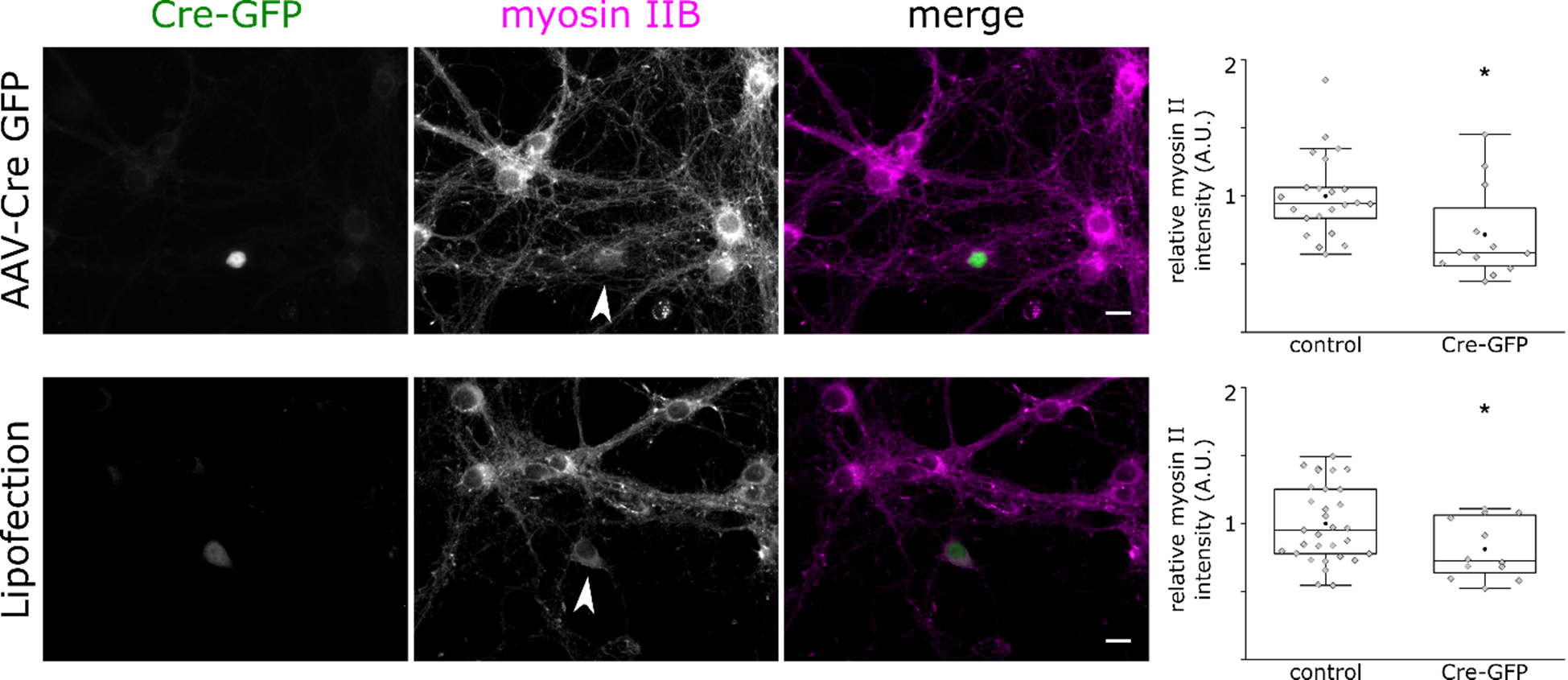
Loss of Tpm3.1 leads to a reduction in myosin IIB immunofluorescence. Cultured hippocampal neurons of conditional *Tpm3* knockout mice (Tp9 line). Arrowheads indicate neurons expressing Cre-GFP after either viral transduction (top panel) or lipofection (bottom panel). We used anti-myosin IIB to compare the distribution of myosin IIB in neurons expressing Cre-GFP and neighboring control neurons. Cre-GFP expressing neurons showed a lower intensity of myosin IIB immunofluorescence (*t* test). Box borders represent the 25^th^ and 75^th^ percentiles, whiskers represent minimum and maximum values less than 1.5x the interquartile range lower or higher than the 25^th^ or 75^th^ percentiles, respectively (Tukey style). Neurons expressing Cre-GFP; transduced: n = 12, 3 independent experiment; transfected n = 12, 3 independent experiments. Control neurons: transduced: n = 22; transfected: n = 31. * denotes statistical significance. *: p < 0.05. Scale bar: 10 µm.

## DISCUSSION

The data presented here show that the tropomyosin isoform Tpm3.1 localizes to the AIS where it decorates actin filaments. Tpm3.1 co-localizes with patches of actin in the AIS that show a low rate of depolymerization. Tpm3.1 shows a periodic distribution but was only partially congruent with sub-membranous actin rings. However, the inhibition of Tpm3.1 affects the alignment of actin rings and the uniformity of their periodicity. In addition, the perturbation of Tpm3.1 function led to the loss of accumulation of ankyrin G and other AIS structural and functional protein, disruption in sorting somatodendritic and axonal proteins, and a reduction in firing frequency. Although an intact actin cytoskeleton is required for the formation of the AIS (Xu and Shrager, 2005), the mature AIS is remarkably stable and insensitive to actin-disrupting drugs (Jones et al., 2014; Leterrier et al., 2015; Sanchez-Ponce et al., 2011; Song et al., 2009). This may lead to the conclusion that the actin cytoskeleton has no significant role in the maintenance of AIS structure. The loss of ankyrin G accumulation upon the perturbation of Tpm3.1 function (Figs. 4 and 5), however, suggests otherwise.

The observed loss of ankyrin G accumulation at the AIS was possibly a consequence of the loss of a latrunculin-resistant population of actin filaments. The periodicity of actin in the AIS is resistant to latrunculin treatment (Abouelezz et al., 2019; Leterrier et al., 2015; Qu et al., 2017). Latrunculin disrupts actin filaments by sequestering monomers, inhibiting polymerization, and favoring depolymerization. Accordingly, filaments with a lower rate of depolymerization are less susceptible to latrunculin treatment compared to dynamic filaments. The negative effect of Tpm3.1 on the depolymerization rate of actin filaments could render AIS actin filaments partially resistant to actin-disrupting drugs like latrunculin (Bach et al., 2009). Indeed, we found that actin filaments in AIS actin patches have a relatively slow rate of depolymerization (Fig. 2). Furthermore, platinum replica electron microscopy showed that the AIS contains short stable actin filaments (Jones et al., 2014). The presence of Tpm3.1 in the AIS and its importance in maintaining AIS structure suggest that it is possible that these filaments are decorated by the 33-34 nm-long Tpm3.1 (Fowler, 1990).

Tpm3.1 enhances the phosphorylation of actin-depolymerizing factor/cofilin (Bryce et al., 2003), thus inhibiting filament severing as well as depolymerization at the pointed ends (Broschat, 1990). Furthermore, Tpm3.1 recruits tropomodulin to the pointed ends (Sung and Lin, 1994), further lowering the rate of depolymerization (Weber et al., 1994; Yamashiro et al., 2014). Thus, the inhibition of Tpm3.1 renders Tpm3.1-decorated actin filaments vulnerable to depolymerization (Bonello et al., 2016) and, therefore, it is plausible that the loss of AIS structure upon Tpm3.1 inhibition is due to a reduction in the stability of actin filaments. Bach et al. (2009) showed that Tpm3.1 is required for stabilizing actin filaments in the formation and maturation of focal adhesions. Tpm3.1-decorated actin filaments are the least sensitive to latrunculin and Cytochalasin D (Creed et al., 2008; Percival et al., 2000). It is, therefore, expected that inhibiting Tpm3.1 function will have a substantial effect on actin filament dynamics in the AIS. Altering the stability, length, or linearity of the actin filaments building the AIS may then lead to less organized structures (Tojkander et al., 2011).

The AIS actin cytoskeleton is believed to comprise sub-membranous actin rings and actin-rich patches (Papandreou and Leterrier, 2018). While sub-membranous actin rings are not exclusive to the AIS (Bertling and Hotulainen, 2017; D’Este et al., 2015; Han et al., 2017; Leite et al., 2016; Leite and Sousa, 2016; Xu et al., 2013; Zhong et al., 2014), actin patches in the AIS are more numerous and resistant to extraction, compared to more distal axonal patches (Balasanyan et al., 2017). Our data suggest that Tpm3.1 co-localizes with actin patches in the AIS (Fig. 3), and that the inhibition of Tpm3.1 led to a reduction in the number of AIS actin patches. Furthermore, in addition to rings and patches, platinum replica electron microscopy revealed the presence of a dense, fibrillar-globular coat over microtubule bundles containing a number of AIS-specific proteins (Jones et al., 2014). Interestingly, this fibrillar coat is specific to the ankyrin G-positive AIS area. However, the thin actin filaments in this fibrillar coat are probably difficult to detect using light microscopy techniques; the coat is likely to altogether look like a faint background staining, with no distinguishable structures. As the distribution of Tpm3.1 was not congruent with sub-membranous actin rings, we propose that Tpm3.1 decorates actin filaments in this fibrillar coat. We expect this fibrillar coat to provide support for the AIS structural complex, which can partially explain the relative loss of uniformity of the sub-membranous actin rings upon Tpm3.1 inhibition. The inhibition of Tpm3.1 may also affect the recruitment and accumulation of other actin-binding proteins that are important for maintaining actin rings or other actin structures in the AIS. In addition to its direct effects on actin filaments, Tpm3.1 also recruits and activates myosin II (Bryce et al., 2003; Gateva et al., 2017), which has recently emerged as an important part of AIS structure (Berger et al., 2018; Evans et al., 2017). It is thus plausible that Tpm3.1 further contributes to the structure of the AIS by recruiting myosin II to the fibrillar coat, providing the lattice with contractile characteristics. This is supported by the relatively low myosin II immunofluorescence in *Tpm3* KO neurons (Fig. 10).

Alternatively, the loss of AIS structure upon Tpm3.1 inhibition may be the result of the perturbation of the specific interactions between Tpm3.1 and proteins contributing to the structure of the AIS; for example, by regulating the binding of βIV-spectrin to the sub-membranous actin rings. However, as ankyrin G accumulation is hardly affected by the depletion of other AIS components within this time-frame (Leterrier, 2016), this scenario is doubtful. In fact, the effect of Tpm3.1 inhibition on firing frequency— likely due to reduced clustering of voltage-gated sodium channels—was relatively rapid (Fig. 8).

Taken together, we believe we can safely say that the AIS contains highly stable Tpm3.1-decorated actin filaments that are essential for AIS structure. These filaments allow the formation of a contractile actin network under the AIS membrane that provides a scaffold for membrane organization and AIS proteins.

## METHODS

### Neuronal cultures, transfections, and preparation of fixed samples

Neuronal cultures were prepared as described previously (Hotulainen et al., 2009). We collected brains from embryonic day 17 Wistar rat fetuses of either sex, removed the meninges, dissected the hippocampi, and dissociated cells in 0.05% papain. Next, we mechanically triturated and suspended the cells in Ca^2+^- and Mg^2+^-free HBBS medium containing sodium pyruvate (1 mM), HEPES (10 mM, pH 7.2), and DNase I (20 U/ml, Sigma-Aldrich). We plated the cells in 24-well plates on glass coverslips 13-mm in diameter coated using poly-L-lysine (0.01 mg/ml, Sigma-Aldrich) in Neurobasal medium (Invitrogen) supplemented with B-27 (Invitrogen), L-glutamine (Invitrogen), and primocin (InvivoGen). For SIM experiments, we used square high-performance glass coverslips 18 x 18 mm (Zeiss) in 6-well cell culture plates. For STORM experiments, we used glass-bottomed 35 mm dishes (Mattek). For experiments involving transfections, we plated neurons at a density of 75,000 cells/cm^2^. For all other experiments, we plated 3,500 cells/cm^2^. We cultured the neurons in a humidified incubator at 37°C and 5% CO_2_, and refreshed the media twice weekly at regular intervals. We used Lipofectamine 2000 (Invitrogen) to transfect 8-10 DIV neurons as described previously (Hotulainen et al., 2009). We used 4% PFA in PBS for 10-15 minutes at room temperature to fix neurons in preparation for immunofluorescence. For the extraction of soluble proteins (Supplementary Figure 5) we treated neurons using 0.1% Triton-X in cytoskeleton buffer (10 mM MES, 150 mM NaCl, 5 mM EGTA, 5 mM glucose, 5 mM MgCl_2_, pH: 6.1) for 60 seconds at room temperature, followed by 4% PFA. For panNa_v_ staining, we used 1% PFA + 3% sucrose in PBS. For EB1 staining, we treated fixed neurons using 1 mM EGTA in 100% methanol for 5 minutes at −20°C, followed by 4% PFA for 5 minutes at room temperature.

We used 0.5% Triton-X in PBS for 10 minutes to permeabilize the cells and 0.2% BSA in PBS (BSA-PBS) for blocking and washing. Following primary and secondary staining, we washed the cells once in 0.1% Triton-X in PBS and 3 times in BSA-PBS. Primary and secondary antibodies were incubated in BSA-PBS at room temperature for 75 minutes and 45 minutes, respectively. For SIM experiments, we incubated γ/9d overnight at 4°C. We mounted coverslips on glass slides using Immu-Mount (Thermo/Shandon). For SIM experiments, we mounted coverslips on glass slides using ProLong Gold (ThermoFisher). Phalloidin was incubated in PBS for 2 hours at room temperature. For STORM experiments, phalloidin staining was performed immediately before imaging.

For AIS intensity experiments using Tp9, Tp16 and C57Bl6 mouse lines, hippocampal neurons were prepared as previously described (Fath et al., 2009). In brief, hippocampi were dissected from the brains of embryonic mice at embryonic day 16 (E16) and dissociated by mechanical trituration after enzymatic exposure to trypsin and DNase I (Sigma-Aldrich). Cells were plated at a density of 58,000 cells/cm^2^ on PDL-coated 12 mm coverslips and maintained in neurobasal media supplemented with 2% B27 and 2 mM Glutamax (Life Technologies) at 37°C and 5% CO_2_. Tp9 cultures were transduced at 0 DIV with either CMV-EGFP-Cre or CMV-EGFP adeno-associated viruses (UNC Vector Core Facility) at a concentration of 5 × 10^7^ viral particles/70,000 cells. Prior to immunostaining, cultures were fixed at 9 DIV with 4% PFA in PBS for 15 min at room temperature. We then permeabilized the cells using 0.1% TritonX-100 in PBS for 5 min and blocked in 2% FBS in PBS. For Tm4 staining, cells were instead permeabilized for 5 min with ice-cold MeOH and blocked in 2% FBS in PBS. Coverslips were mounted on glass slides using ProLong Gold (Life Technologies).

### Plasmids, antibodies and reagents

mCherry-C1 (mCherry) was purchased from Clontech. PAGFP-actin (Dopie et al., 2012) was a kind gift from Maria Vartiainen (University of Helsinki, Finland). YFP-Tpm3.1 and YFP-Tpm3.2 were described previously (Tojkander et al., 2011). shRNA against rat Tpm3.1 (NM_173111.1) was purchased from GeneCopoeia (RSH053175-33-mH1, target sequence CCAAGTCTTAGCCAAACAACA). Mouse monoclonal anti-ankyrin G antibody (1:1000, UC Davis/NIH NeuroMab Facility, Clone 106/36) and mouse monoclonal panNF-186 (1:500, UC Davis/NIH NeuroMab Facility, Clone A12/18) were purchased from NeuroMab. Sheep polyclonal anti-γ/9d (1:100, for SIM: 1:50, AB5447), mouse monoclonal anti-Tpm3 (clone 2G10.2, MAB2256), rabbit polyclonal anti-GluA1 (1:300, Chemicon/Millipore AB1504), chicken polyclonal anti-MAP2 (1:10000, AB5543), chicken anti-β3 tubulin (1:500, AB9354) and mouse anti-ankyrin G (1:500, MABN466) were purchased from Merck Millipore. Rabbit polyclonal anti-panNa_v_ was purchased from Alomone Labs (1:200, ASC-003). Rabbit anti-myosin IIB (1:500) was purchased from Australian Biosearch (909901). Rabbit anti-TRIM46 (1:100) was described previously (van Beuningen et al., 2015). Mouse anti-EB1 (1:100, 610535) were purchased from BD Transduction Lab. Rabbit anti-GFP (1:1000, AB290) was purchased from Abcam. Rabbit polyclonal δ9d (1:250, purified serum) was produced by the Gunning lab and previously described (Schevzov et al., 2011). Alexa 647- and Alexa 488-conjugated phalloidin were purchased from ThermoFisher (1 μM, A22287). Alexa Fluor-conjugated secondary antibodies (1:400) were purchased from ThermoFisher. The anti-tropomyosin drugs TR100 and Anisina (ATM3507) were described previously (Currier et al., 2017; Stehn et al., 2016; Stehn et al., 2013) and were added to culture media from 50-mM stock solutions in DMSO. Latrunculin B (Sigma, L5288) was added to culture media from a 2.5-mM stock solution in DMSO.

### Generation of the Tpm3 gene exon 1b conditional knockout mouse

Genomic fragments of the *Tpm3* exon 1b region were assembled into a targeting vector containing a Neo cassette flanked by Frt sites located upstream of exon 1b (Supplementary Figure 9). LoxP sites were inserted upstream of the Neo cassette and in the 3’ region downstream from exon 1b. A diphtheria toxin A (DTA) cassette was used in the targeting construct for negative selection. ES cells were electroporated with the targeting construct and the selected ES cell clones were microinjected into C57Bl/6 blastocysts to generate chimeras containing the targeted allele. Chimeric mice were subsequently bred against Flp transgenic mice to remove the Neo drug selection cassette via Flp-mediated recombination and obtain germline F1 mutants with the conditional knockout allele. Exon 1b can be excised to create the constitutive full knockout allele by Cre-mediated recombination of the LoxP sites in the conditional knockout allele. The conditional knockout allele prior to Cre-mediated recombination was tested by PCR genotyping (Supplementary Figure 9b).

### Imaging

For taking confocal stacks, we used a Zeiss LSM880 inverted confocal microscope (Zeiss) equipped with a 63x 1.40 NA oil-immersion objective or a Zeiss LSM710 upright confocal microscope (Zeiss) equipped with a 63x 1.46 NA oil-immersion objective. Both microscopes were equipped with 405, 488, 561, and 633 nm laser lines. We adjusted laser power and gain settings as to maximize signal-to-noise ratio while ensuring no pixels were saturated outside the somata. *Z* stacks were acquired at a step size of 0.2 μm. Imaging of fixed samples was performed at room temperature. For live-cell imaging we used a temperature-controlled chamber and CO_2_ supply. We imaged live cells in culture media at 37°C and 5% CO_2_ using a Zeiss LSM710 upright confocal microscope equipped with a 63x 1.0 NA water-dipping objective. For epifluorescence imaging, we used a Zeiss Axio Imager Z2 upright epifluorescence microscope (Zeiss) equipped with a 40x 1.3 NA oil-immersion objective and a Hamamatsu Orca Flash 4.0 LT camera (Hamamatsu). Alternatively, we used a Zeiss Axio Imager M1 upright epifluorescence microscope (Zeiss) equipped with a 20x 0.8 NA objective and an AxioCam HRm camera (Zeiss). We used Zeiss Zen software for acquisition.

For SIM, we used a DeltaVision OMX SR imaging system (GE Healthcare Life Sciences) equipped with a 60x 1.42 NA PlanApo N oil objective, 488, 560, and 640 nm laser lines, and 3 sCOMS cameras. We used the AcquireSR software for acquisition and SoftWorx software for image reconstruction and alignment.

For STORM, we used an N-STORM system comprising a Nikon Eclipse Ti-E inverted microscope with Nikon IR-based Perfect Focus System (Nikon Instruments). The microscope was equipped with a 100x 1.49 NA Apo TIRF oil-immersion objective, 405 and 647 nm laser lines (100 and 300 mW, respectively), and an iXon+ 897 camera (Andor). We used a Nikon Intensilight metal arc light source (Nikon) to locate neurons. We used NIS-Elements software for acquisition. We placed the neurons in STORM buffer immediately before imaging: pH 7.9 50 mM Tris, 10% glucose, 70 mM freshly prepared MEA (Sigma), 0.75 mg/ml glucose oxidase (Sigma), and 0.04 mg/ml catalase (Sigma). We continuously illuminated the sample using the 647-nm laser at full power for a series of 20,000-30,000 images of 20-40 ms exposure time (128 x 128 pixels). We used an increasing intensity of 405-nm illumination to reactivate fluorophores. For the analysis of the AIS in mouse hippocampal neurons, fixed samples were imaged on an Axioskop 40 microscope (Zeiss) using a 40x oil objective. AIS intensity was measured using ImageJ software and statistically analyzed in GraphPad Prism (v7.02).

### Photoactivation

For photoactivation experiments, live neurons were stained using panNF-186 to label the AISs (Hedstrom et al., 2008). 1-2 hours before imaging, neurons were incubated in culture media containing panNF-186 for 10 minutes at 37°C and 5% CO_2_, washed 3 times in Neurobasal media, then incubated in culture media containing anti-mouse Alexa Fluor-647 for 10 minutes at 37°C and 5% CO_2_. Neurons were then washed and returned to culture media. We acquired a frame every 3 seconds covering an area of 38.4 x 38.4 μm (256 x 256 pixels) with a pixel dwell time of 3.15 μs. After acquiring 3 pre-activation frames, we used 10-15 iterations of a 405-nm laser at full power (30 mW) to induce photoactivation. To measure the rate of fluorescence decay we limited the photoactivation to a square region of interest 2.25 x 2.25 μm (15 x 15 pixels). Acquisition was resumed immediately after photoactivation for 360 s at 3-s intervals.

For each experiment, the fluorescence intensity within the region of interest in the pre-activation frames was measured, averaged, and the value was subtracted from subsequent measurements. The first post-activation frame was taken as time-point 0 s. The average fluorescence intensity in subsequent frames (*F*_*t*_) were normalized to the first frame (*F*_*0*_) to plot fluorescence decay curves. Decay curves were fit to the two-component exponential function 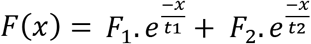 where *x* is time, *F*_1_ is the component with the smaller time constant (dynamic pool), *F*_2_ is the component with the larger time constant (stable pool), and *t*_1_ and *t*_2_ are the respective time constants (Star et al., 2002). The proportions of the dynamic and stable pools were calculated as 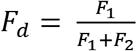 and 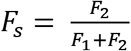, respectively.

### Electrophysiology

For electrophysiological recordings, we placed coverslips in a submerged recording chamber and perfused using an extracellular solution containing (in mM): 124 NaCl, 3 KCl, 1.25 NaH_2_PO_4_, 1 MgSO_4_, 26 NaHCO_3_, 15 D-glucose, 2 CaCl_2_; bubbled using 5% CO_2_/95% O_2_ at 32°C. We used patch electrodes (3-5 MΩ) to perform whole-cell recordings in individual cells. Whole-cell pipettes used for current-clamp experiments contained the following (in mM): 130 K-gluconate, 8 NaCl, 10 HEPES, 0.4 EGTA, 4 Mg-ATP and 0.3 Na-GTP, with the addition of either Anisina (2.5 µM) or DMSO (0.2%). The osmolarity of all intracellular solutions was adjusted to 285 mOsm and the pH to 7.25. Action potentials were detected and analyzed using the Mini Analysis Program 5.6.6. (Synaptosoft). Firing frequency was calculated from the interval between the second and the third action potential during a 500 ms depolarizing step evoked by 100-200 pA. Data are expressed as frequency.

### Image Analysis

We used the Fiji software platform for image analysis (Schindelin et al., 2012). We plotted the fluorescence intensity profile ~1-5 μm along the AIS where periodicity was visible in a single plane in a SIM reconstruction, avoiding patches and fasciculations. We used a MATLAB script to locally normalize fluorescence intensity in each profile. We used the “autocorr” function in MATLAB to calculate the autocorrelation function for each profile and obtain autocorrelation curves. We used the “findpeaks” function in MATLAB to detect individual peaks in each profile and note their locations to calculate inter-peak distances.

To quantify the effect of Tpm3.1/2 knockdown on the accumulation of ankyrin G at the AIS, we used NeuronJ (Meijering et al., 2004) to blindly trace the AISs of transfected neurons expressing either scramble or anti-Tpm3.1/2 shRNA, as well as neighboring non-transfected neurons. We used a 3-μm moving average to smooth fluorescence intensity profiles (van Beuningen et al., 2015) and noted the peak intensity of every profile. Relative ankyrin G fluorescence intensity was calculated by comparing the peak intensity recorded in the transfected neuron to the mean peak intensity of the neighboring non-transfected neurons. We then normalized the mean values to the mean relative intensity of neurons expressing scramble shRNA.

For calculating the ALI, we measured the fluorescence intensity profile of either anti-ankyrin G or panNa_v_ in the initial 30 μm of every neurite in a single neuron. After subtracting background fluorescence, we calculated the 3-μm moving average to smooth the profiles (van Beuningen et al., 2015), trimmed the profiles for even averaging, and then normalized all measurements to the peak intensity value. We then noted the normalized peak value for each neurite and used them to calculate the median peak value. AIS localization index was expressed as *ALI* = 1 − *median*_*peaks*_ such that a value closer to 1 indicates the strongest accumulation in a single neurite, while a value closer to 0 indicates homogenous concentration across all neurites.

For calculating the ADR, we used NeuronJ (Meijering et al., 2004) to trace the axon and dendrites in confocal stacks. We then used a maximum intensity projection to measure background fluorescence, the mean fluorescence intensity along ~100 μm of the axon distal to the AIS, and the mean fluorescence along all dendrites. ADR was calculated as 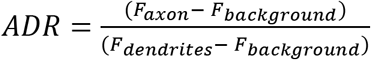 (Lewis et al., 2009).

We used custom scripts written by Leterrier et al. (2015) to convert STORM localization files into a format readable by the ThunderSTORM plugin (Ovesny et al., 2014), which was used for reconstructing images. For evaluating periodicity, we used MATLAB to fit intensity profiles to a single-term Fourier series represented by *y* = *a*_0_ + *a*_*i*_ cos(*iwx*) + *b*_*i*_ sin(*iwx*) and obtained the period length and the R-squared value reflecting goodness of fit.

### Statistical Analyses

Binomial tests were performed using the SPSS software package. All other statistical analyses were performed using the OriginLab software package. Intensity profiles curves for STORM data were created using MATLAB. All other graphs were created using OriginLab. Data were checked for normality (Shapiro-Wilk test) and homogeneity of variance (Levene’s test). For parametric data, we used the independent two-sample *t* test, or one-way ANOVA and Tukey’s test for post-hoc analysis. For non-parametric data, we used Kruskal-Wallis ANOVA and Mann-Whitney *U* Test. We used Kolmogorov-Smirnov test to compare the distributions of inter-peak distances under different conditions. We created box plots in Tukey style (Krzywinski and Altman, 2014) with the addition of solid black circles to represent the mean. Classification of photoactivation data as belonging to actin patches or areas outside the patches was done using hierarchical cluster analysis. We used double-exponential decay fits of 44 normalized fluorescence decay curves recorded from the AIS, each responding to 120 variables corresponding to fluorescence levels at 120 time points between 0 and 357. This number of time points corresponds to the sampling frequency of the recordings. We ran a hierarchical cluster analysis using Ward’s method (squared Euclidean distance, data standardization: range 0-1) that assigned 15 curves as belonging to AIS actin patches, and 29 curves as recorded from areas outside the patches. To test the significance of the difference between the resultant clusters, we performed one-way ANOVA for each time point. The between-groups difference was significant at all time points after 0 s (p < 0.05). The difference between recordings from dendrites and areas in the AIS outside the actin patches was not significant at any time point (Tukey’s test), while recordings from the patches were significantly different from recordings from both dendrites and areas outside the patches (Tukey’s test, p < 0.05) for all time points after 0 s.

## ACKNOWLEDGMENTS

We thank Seija Lågas, Outi Nikkilä, and Iryna Hlushchenko for their help with neuronal culture preparation. We are grateful to Veijo Salo for his assistance with the STORM experiments and buffer preparations. We are grateful to Sari Lauri for generously providing anti-GluA1. We are grateful to Enni Bertling for her critical comments on the manuscript. We thank Jeff Hook for his help with the generation of the conditional exon 1b knockout mouse, and Nicole Bryce for preparation of the Supplementary Figure 9. All imaging was performed using microscopes in the Neuroscience Center and Biomedicum Imaging Unit.

This work was supported by the Academy of Finland (PH, SA 266351) and Doctoral Programme Brain & Mind (AA), the Australian Research Council (DP180101473) (TF), the Australian National Health and Medical Research Council (APP1083209) (TF and PWG), (APP1079866, APP1100202) (ECH and PWG) and The Kids Cancer Project (ECH and PWG).

The authors declare no competing financial interests except for PWG and ECH who own shares in a company developing anti-tropomyosin drugs.

## AUTHOR CONTRIBUTIONS

A. Abouelezz, and P. Hotulainen designed experiments with the exception of experiments presented in Fig. 5, 8, and 10, and Supplementary figures 7, 9 and 10. T. Fath designed experiments presented in Fig. 5 and Supplementary Figure 10 and C.C. Hoogenraad designed the experiments presented in Supplementary Figure 7. T. Taira and M. Segerstråle designed the experiments presented in Fig. 8. A. Abouelezz performed all experiments and analyzed all data, with the exception of data and analyses presented in Fig. 5, 8, and 10, and Supplementary figures 9 and 10. H. Stefen and T. Fath performed experiments and analyzed data presented in Fig. 5 and 10, and Supplementary Figure 10. T. Taira and M. Segerstråle performed the experiments and analyzed data presented Fig. 8. E. Hardeman provided conditional *Tpm3 k*nockout mice and Figure S9. D. Micinski assisted with the analysis of the experiments presented in Supplementary figures 11 and 12. R. Minkeviciene assisted with the preparation of neuronal cultures for the experiments presented in Fig. 8. P.W. Gunning provided tropomyosin constructs, antibody, and inhibitors. C.C. Hoogenraad provided antibodies for TRIM46, EB1 and NF-186. A. Abouelezz and P. Hotulainen prepared figures and wrote and edited the manuscript.

## Notes

#### Summary of Updates

Spelling of author name corrected.

