## Supplementary Figures for "Tropomyosin Tpm3.1 is required to maintain the structure and function of the axon initial segment"

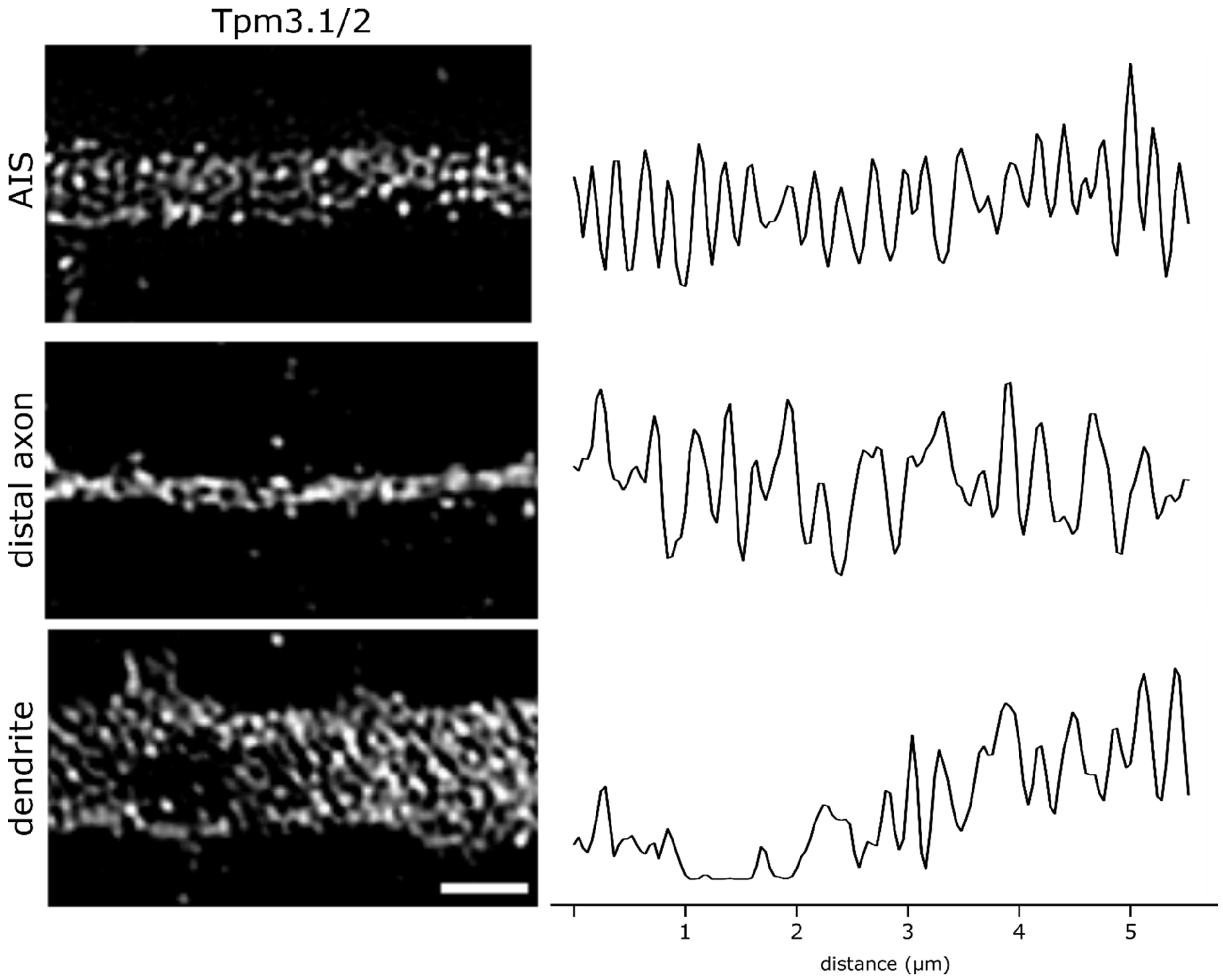


**Supplementary figure 1 | Tpm3.1 does not exhibit clear periodicity in the distal axon or in dendrites of cultured rat hippocampal neurons.** *Left*: SIM reconstructions of the AIS, a distal region of the axon, and a dendrite of a rat hippocampal neuron at 14 DIV stained using anti-γ/9d. Periodic Tpm3.1/2 immunofluorescence is visible in the AIS, but not in the distal regions of the axon or in dendrites. *Right*: Anti-γ/9d fluorescence intensity profiles from the corresponding images. Scale bar: 1 μm.

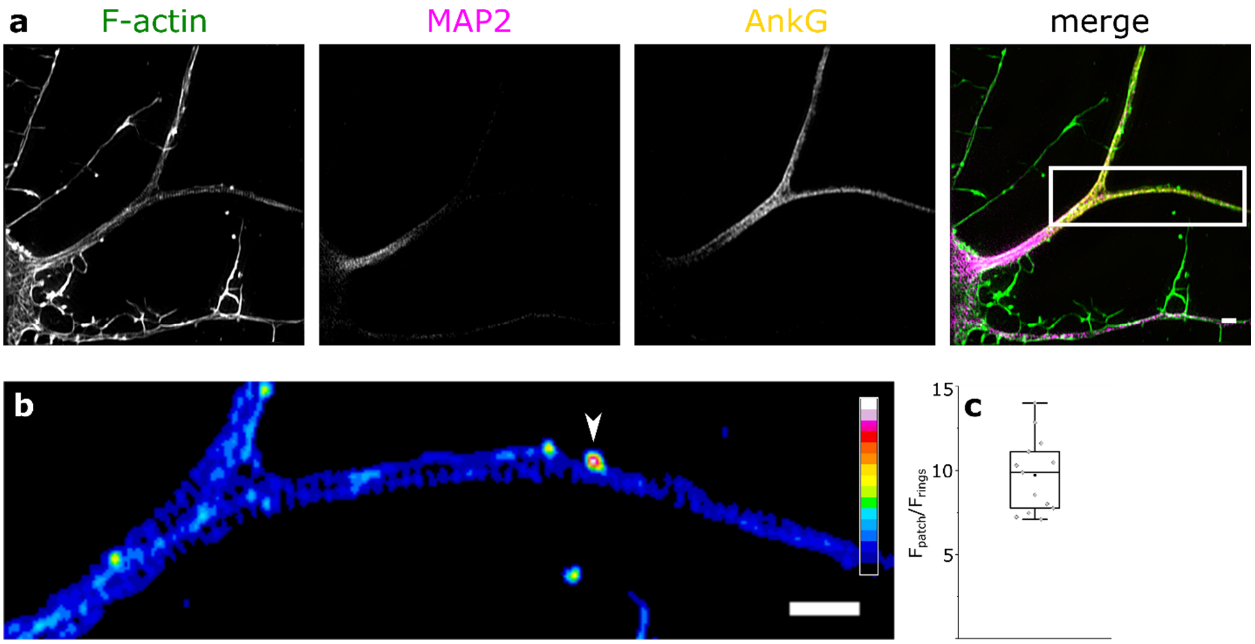


**Supplementary figure 2 | AIS actin patches contain more F-actin than sub-membranous actin rings.** **a** SIM reconstructions of rat hippocampal neuron at 14 DIV stained using Alexa 488-tagged phalloidin, anti-MAP2, and anti-ankyrin G. The boxed region is enlarged in (**b**). **b** Maximum intensity projection of a SIM reconstruction of F-actin in the boxed region, corresponding to the AIS. Color code indicates normalized fluorescence intensity levels. Arrowhead indicates F-actin patch. **c** The median fluorescence intensity of phalloidin in AIS actin patches was, on average, 9.7 times that of sub-membranous actin rings. Black circle represents mean value. Box borders represent the 25$^{th}$ and 75$^{th}$ percentiles, whiskers represent minimum and maximum values less than 1.5x the interquartile range lower or higher than the 25$^{th}$ or 75$^{th}$ percentiles, respectively (Tukey style). n = 13, 4 independent experiments, same data set used in Fig. 9. Scale bar: 1 µm.

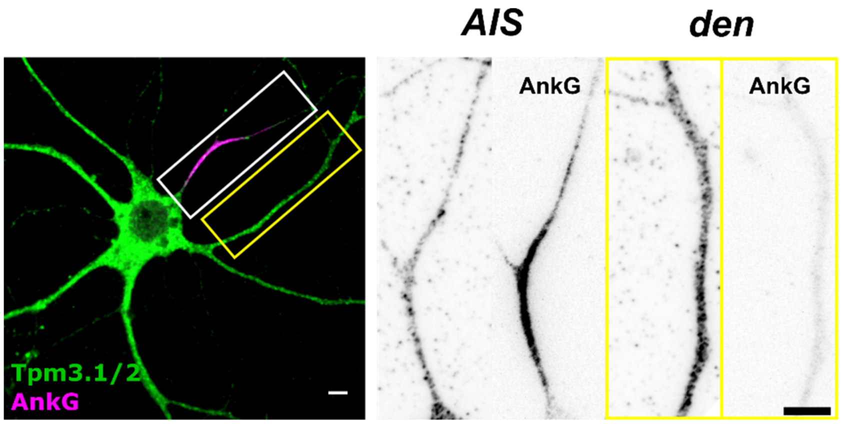


**Supplementary figure 3 | Endogenous Tpm3.1 shows a non-uniform distribution similar to AIS actin patches.** Tpm3.1 was visualized in rat hippocampal neurons at 10 DIV using anti-γ/9d. Anti-ankyrin G served to label the AIS. Anti-γ/9d simultaneously detects Tpm3.1 and Tpm3.2. Anti-γ/9d immunofluorescence in the AIS was unevenly distributed in the AIS, with patches of high intensity similar to AIS actin patches. In contrast, anti-γ/9d in the somatodendritic domain was diffuse and ubiquitous. Scale bar: 5 μm.

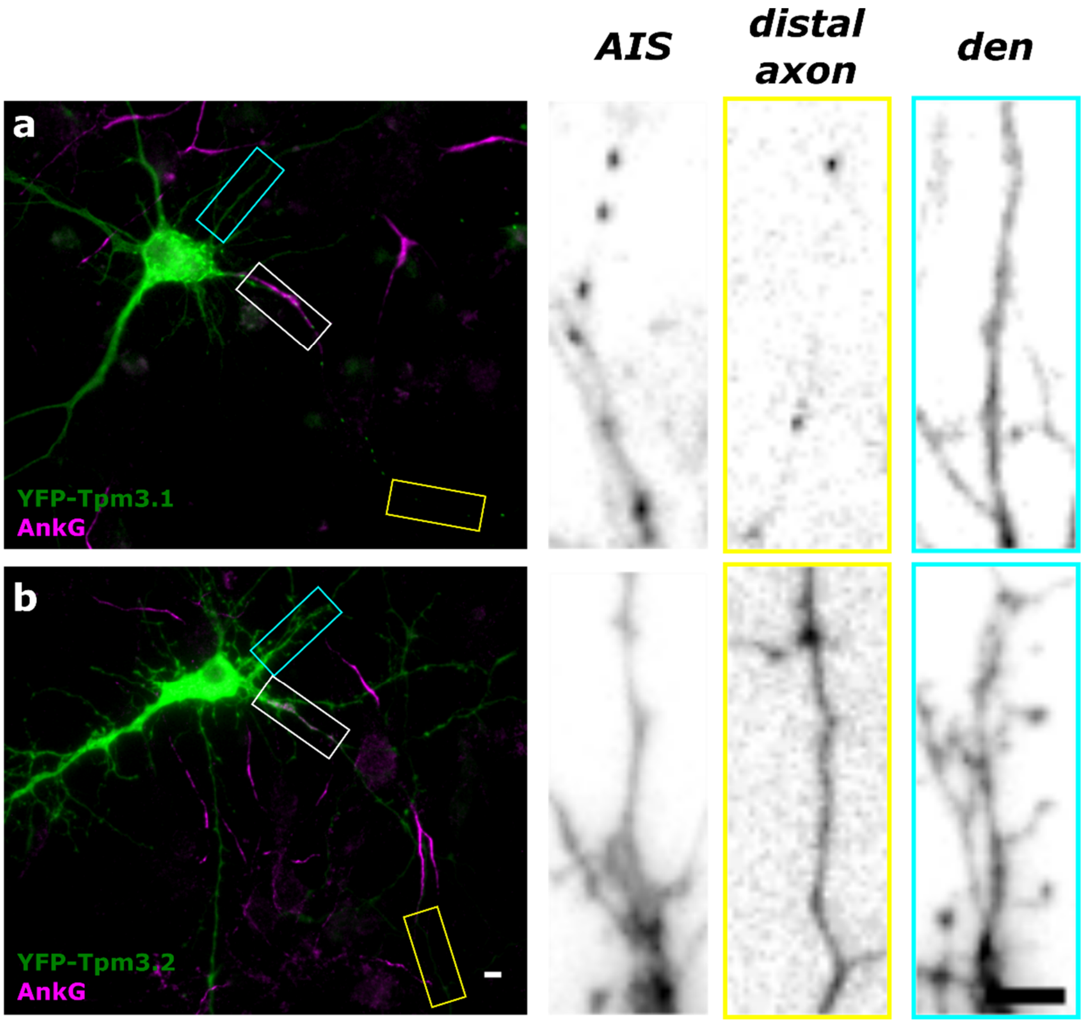


**Supplementary figure 4 | Exogenous tropomyosin isoform Tpm3.1 localizes to the AIS. a** Rat hippocampal neurons expressing either YFP-Tpm3.1 (**a**) or YFP-Tpm3.2 (**b**). Neurons were fixed 8 hours post-transfection. Anti-ankyrin G served to label the AIS. Patches of YFP-Tpm3.1 can be seen in the AIS (white box) and distally in the axon (yellow box) while the somatodendritic domain (cyan box) shows a diffuse, less intense distribution. YFP-Tpm3.2 shows a diffuse staining in the AIS and distal regions of the axon, and a much stronger presence in the somatodendritic compartment and dendritic spines. Scale bar: 10 µm.

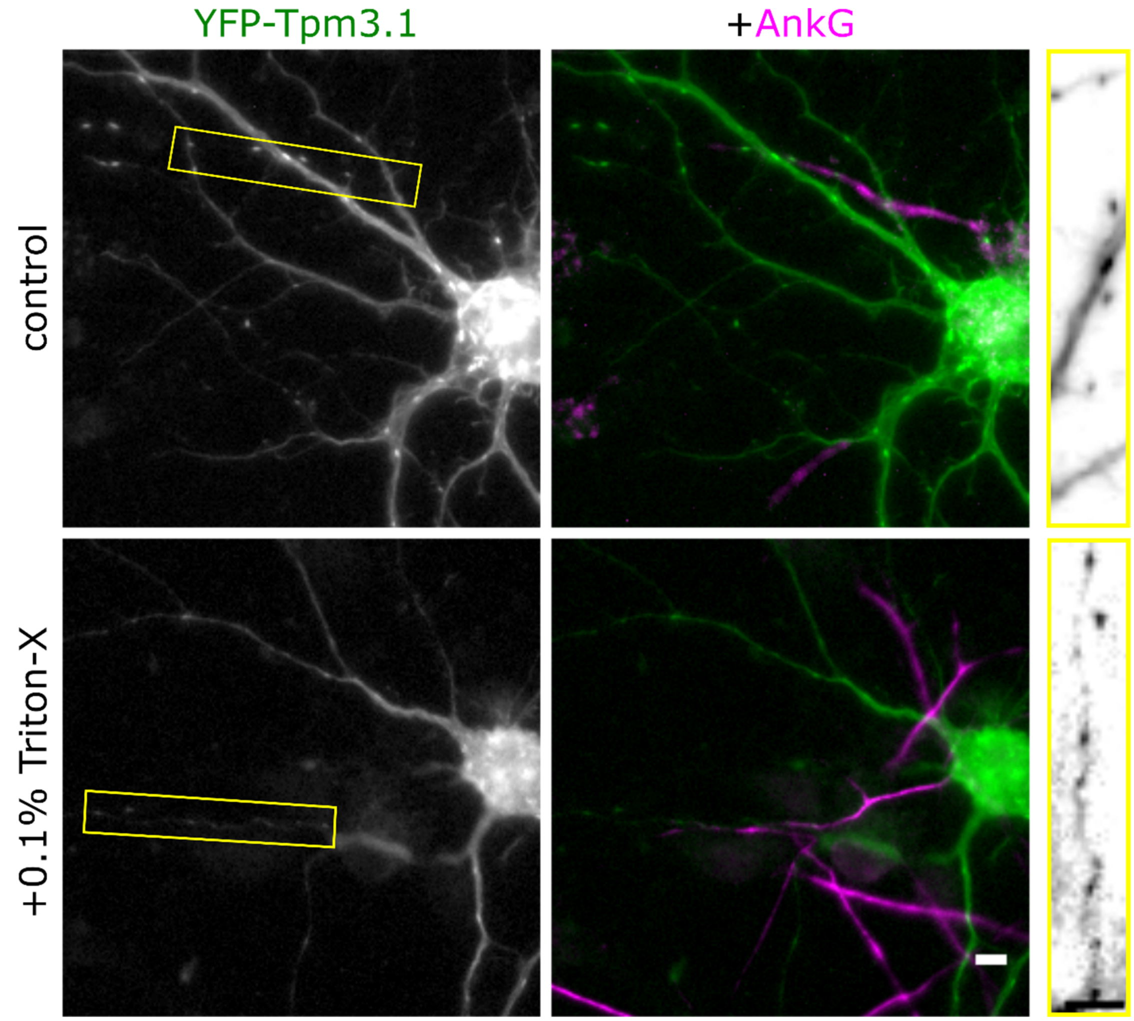


**Supplementary figure 5 | Tropomyosin isoform Tpm3.1 forms bright puncta in the AIS that are resistant to detergent extraction.** Rat hippocampal neurons expressing either YFP-Tpm3.1 were either fixed or extracted in 0.1% Triton-X for 60 seconds then fixed 8 hours post-transfection. Anti-ankyrin G served to label the AIS. Patches of YFP-Tpm3.1 can still be seen in the AIS after detergent extraction. Scale bar: 5 µm.

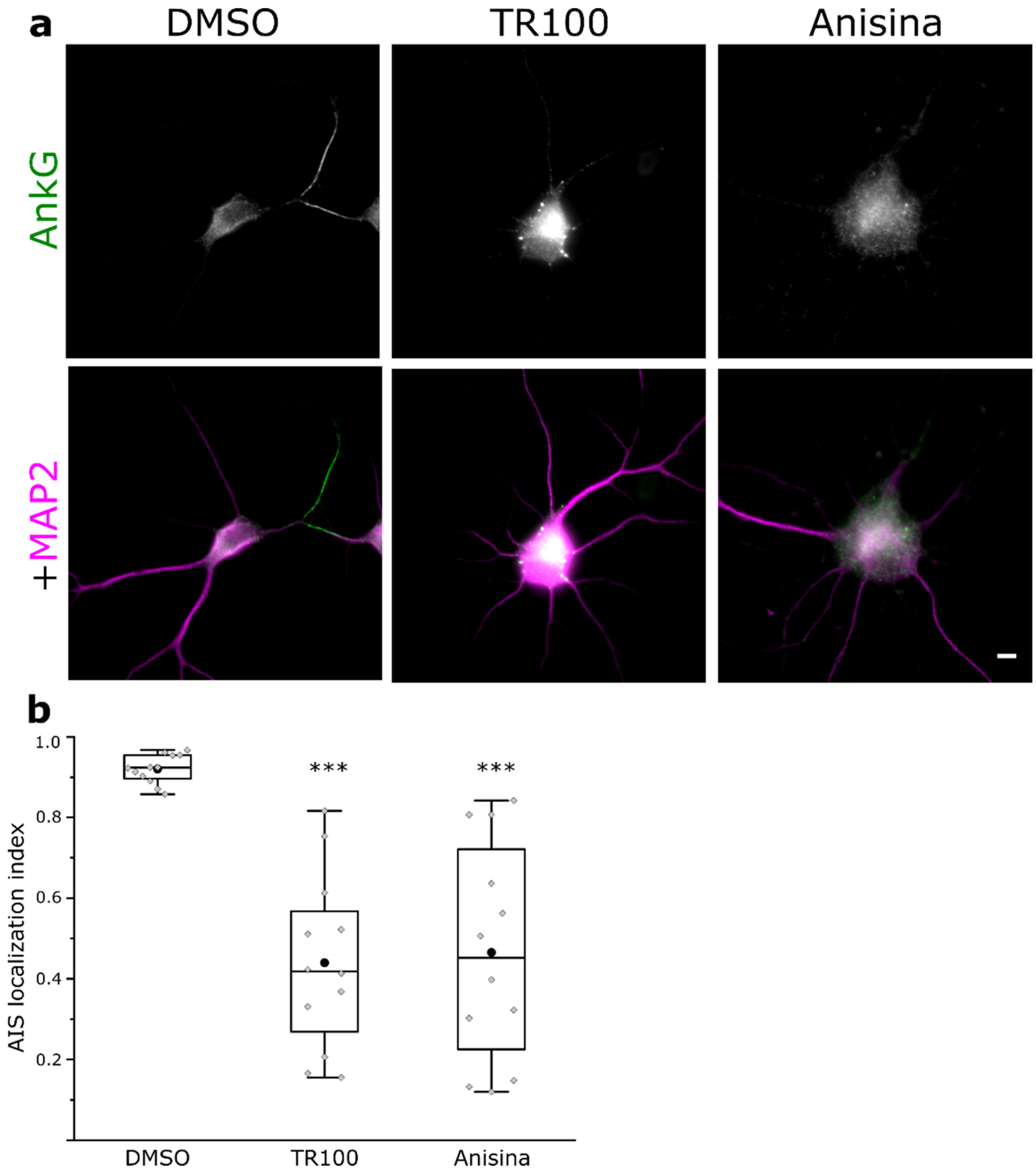


**Supplementary figure 6 | Overnight inhibition of Tpm3.1 reduces the accumulation of ankyrin G at the AIS. a** Rat hippocampal neurons treated overnight at 9 DIV using DMSO or the small-molecule Tpm3.1 inhibitors TR100 or Anisina (ATM3507). Anti-MAP2 served to label the somatodendritic domain, anti-ankyrin G served to measure the accumulation of ankyrin G. **b** AIS localization indices for neurons treated using DMSO, TR100 (5 µM), or Anisina (2.5 µM). Both TR100- (mean ALI: 0.44 ± 0.06, mean ± SEM) and Anisina-treated neurons (mean ALI: 0.48 ± 0.09, mean ± SEM) were significantly different from DMSO controls (mean ALI: 0.92 ± 0.01, mean ± SEM; Mann-Whitney *U* test). Black circles represent mean values. Box borders represent the 25th and 75th percentiles, whiskers represent minimum and maximum values less than 1.5x the interquartile range lower or higher than the 25th or 75th percentiles, respectively (Tukey style). For each treatment: n = 12, 3 independent experiments. Scale bar: 5 µm.

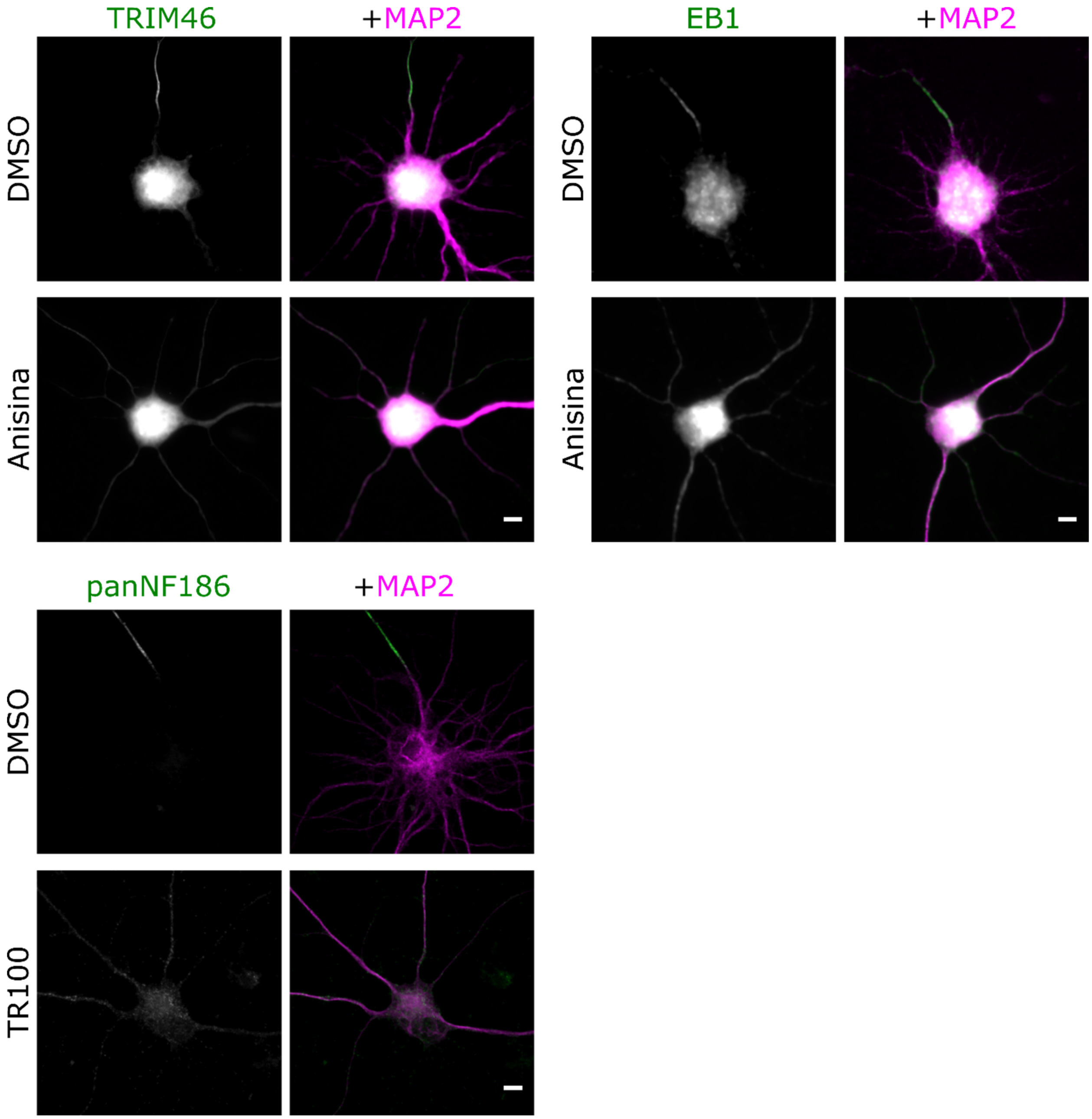


**Supplementary figure 7 | Tpm3.1 inhibition leads to the loss of accumulation of AIS markers TRIM46, EB1, and NF-186 at the AIS.** Rat hippocampal neurons treated at 9-11 DIV using, Anisina (5 µM for 6 hours), TR100 (5 µM overnight), or equivalent volumes of DMSO. Anti-MAP2 served to label the somatodendritic domain. Anti-ankyrin G, anti-TRIM46, anti-EB1, or panNF-186 served to visualize the corresponding AIS marker. DMSO-treated neurons showed typical accumulation of TRIM46 and EB1, which are implicated in the regulation of microtubules, as well as the AIS-specific adhesion protein neurofascin-186 (NF-186). Anisina- or TR100-treated neurons showed no accumulation of any of these proteins at the AIS. Scale bar: 5 µm.

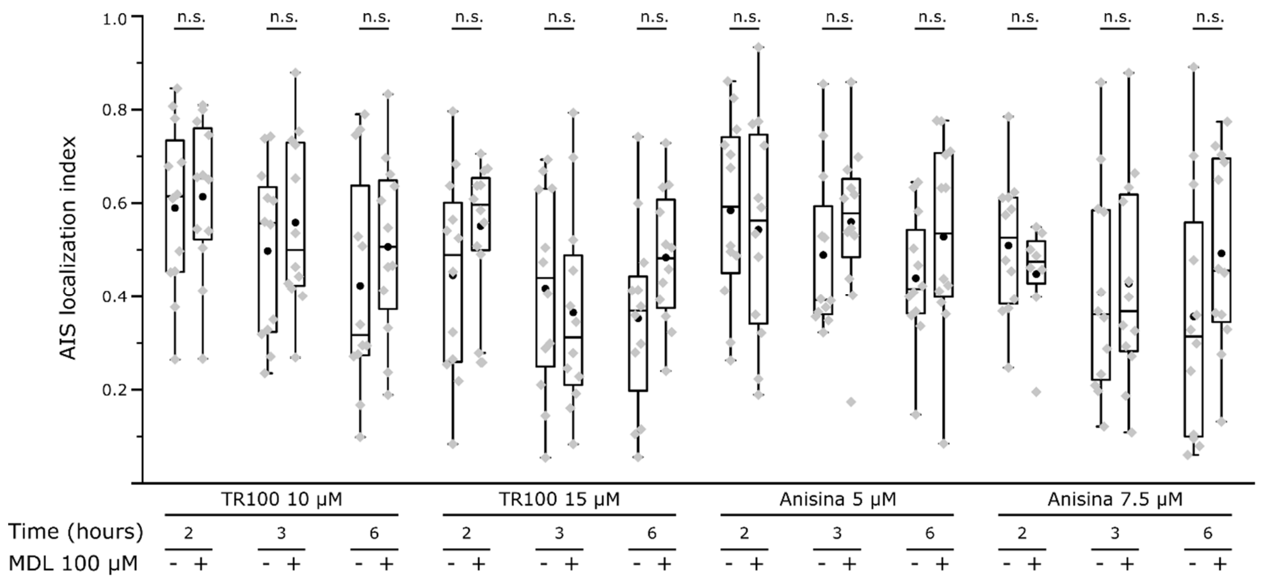


**Supplementary figure 8 | The effect of Tpm3.1 inhibition on the accumulation of ankyrin G at the AIS is calpain-independent.** AIS localization indices (ALI) for ankyrin G in rat hippocampal neurons treated at 10 DIV using the small-molecule Tpm3.1 inhibitors TR100 or Anisina (ATM3507) for 2, 3, or 6 hours, in the presence or absence of the calpain inhibitor MDL28170 (100 μM). The presence of MDL28170 did not significantly affect the ALI for any of the treatments (Mann-Whitney *U* test). Black circles represent mean value. Box borders represent the 25$^{th}$ and 75$^{th}$ percentiles, whiskers represent minimum and maximum values less than 1.5x the interquartile range lower or higher than the 25$^{th}$ or 75$^{th}$ percentiles, respectively (Tukey style). Anisina 7.5 μM + MDL28170 100 μM, 2 hours: n = 8, 2 independent experiments; all other treatments: n = 12, 3 independent experiments. Part of these data were used for Figure 4. n.s.: not significant.

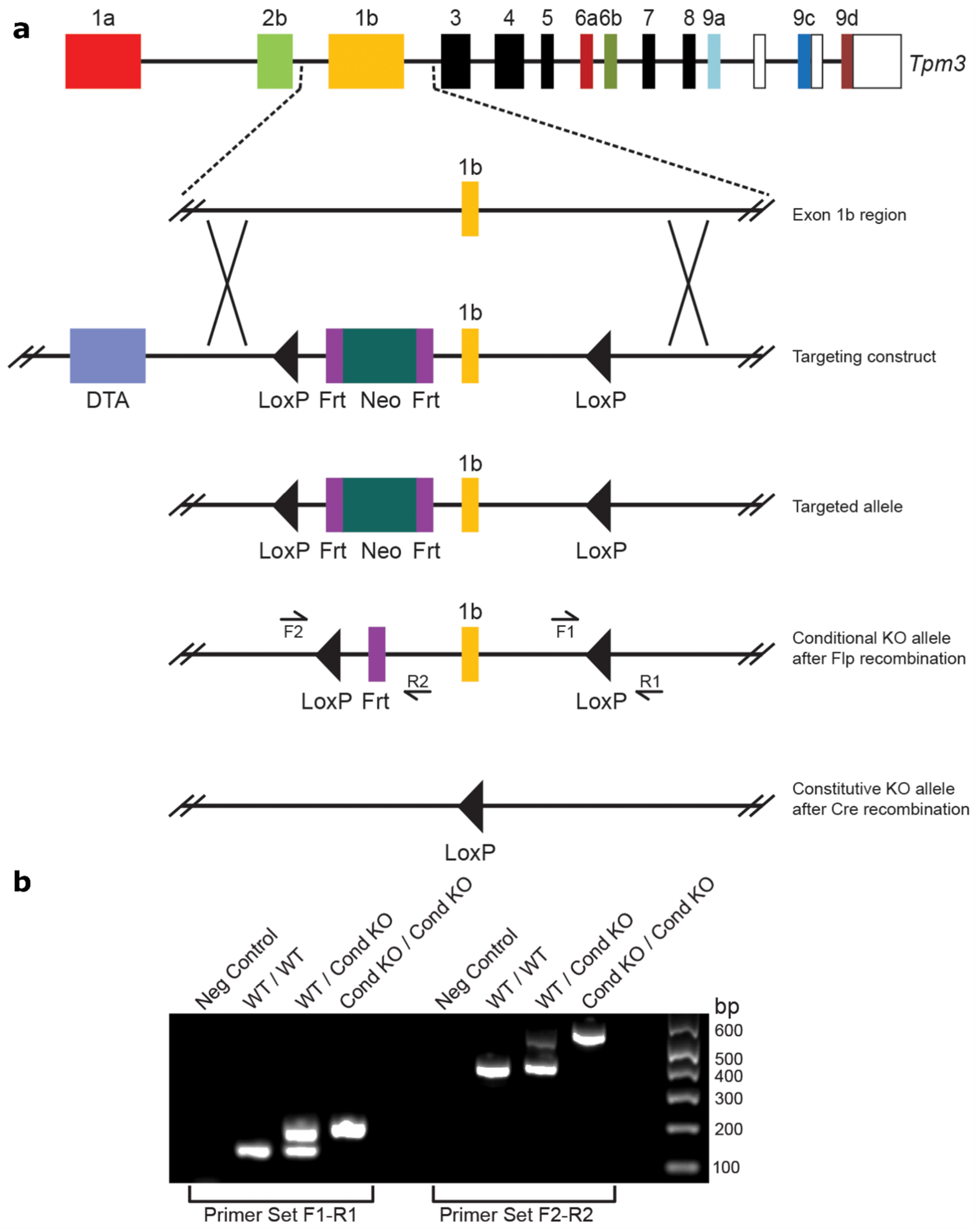


**Supplementary figure 9 | Generation of the Tpm3 gene exon 1b knockout mouse. a** Genomic fragments of the Exon 1b region were assembled into a targeting vector containing a Neo cassette flanked by Frt sites located upstream of exon 1b. LoxP sites were inserted upstream of the Neo cassette and in the 3' region downstream from exon 1b. A diphtheria toxin A (DTA) cassette was used in the targeting construct for negative selection. ES cells were electroporated with the targeting construct and the selected ES cell clones were microinjected into C57Bl/6 blastocysts to generate chimeras containing the targeted allele. Chimeric mice were subsequently bred against Flp transgenic mice to remove the Neo drug selection cassette via Flp-mediated recombination and obtain germline F1 mutants with the conditional knockout allele. The location of the PCR genotyping primer pairs is indicated (arrows, F1-R1 and F2-R2). Exon 1b can be excised to create the constitutive full knockout allele by Cre-mediated recombination of the LoxP sites in the conditional knockout allele. **b** PCR genotyping of the conditional knockout allele prior to Cre-mediated recombination. Primer set 1: The wild type (WT) allele is 151 bp and the conditional knockout (Cond KO) allele is 212 bp. Primer Set 2: The wild type (WT) allele is 402 bp and the conditional knockout (Cond KO) allele is 536 bp. A negative (Neg) control for each primer set containing no template DNA is also shown. For heterozygous mice (WT/CondKO) the bands generated by the F2-R2 primer set favored the smaller sized band in abundance; whereas a more even distribution was seen with the bands generated by the F1-R1 primer set.

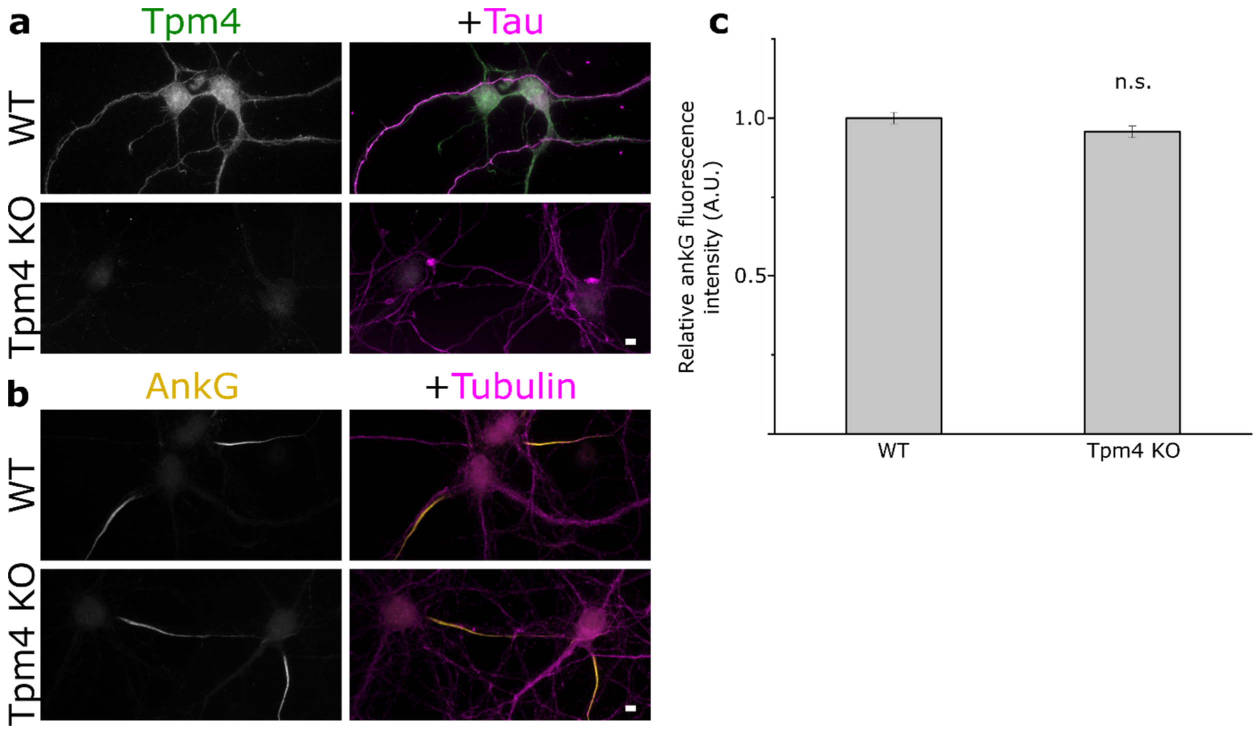


**Supplementary figure 10 | *Tpm4* knockout mice show typical accumulation of ankyrin G at the AIS.** **a** Cultured hippocampal neurons of wild-type and *Tpm4* knockout mice showing the absence of Tpm4 immunofluorescence in the knockouts. Anti-Tau served to label axons. **b** *Tpm4* knockout mice show typical accumulation of ankyrin G. β3-tubulin served to label neurons. **c** The difference in the mean AIS ankyrin G fluorescence intensity between wild-type (1 ± 0.02, mean ± SEM) and knock-out (0.96 ± 0.02, mean ± SEM) neurons was not statistically significant (Mann-Whitney *U* test). Wild-type: n = 122, 3 independent experiments; *Tpm4* knockout: n = 147, 3 independent experiments. Error bars represent standard error of mean. n.s.: not significant. Scale bar: 5 μm.

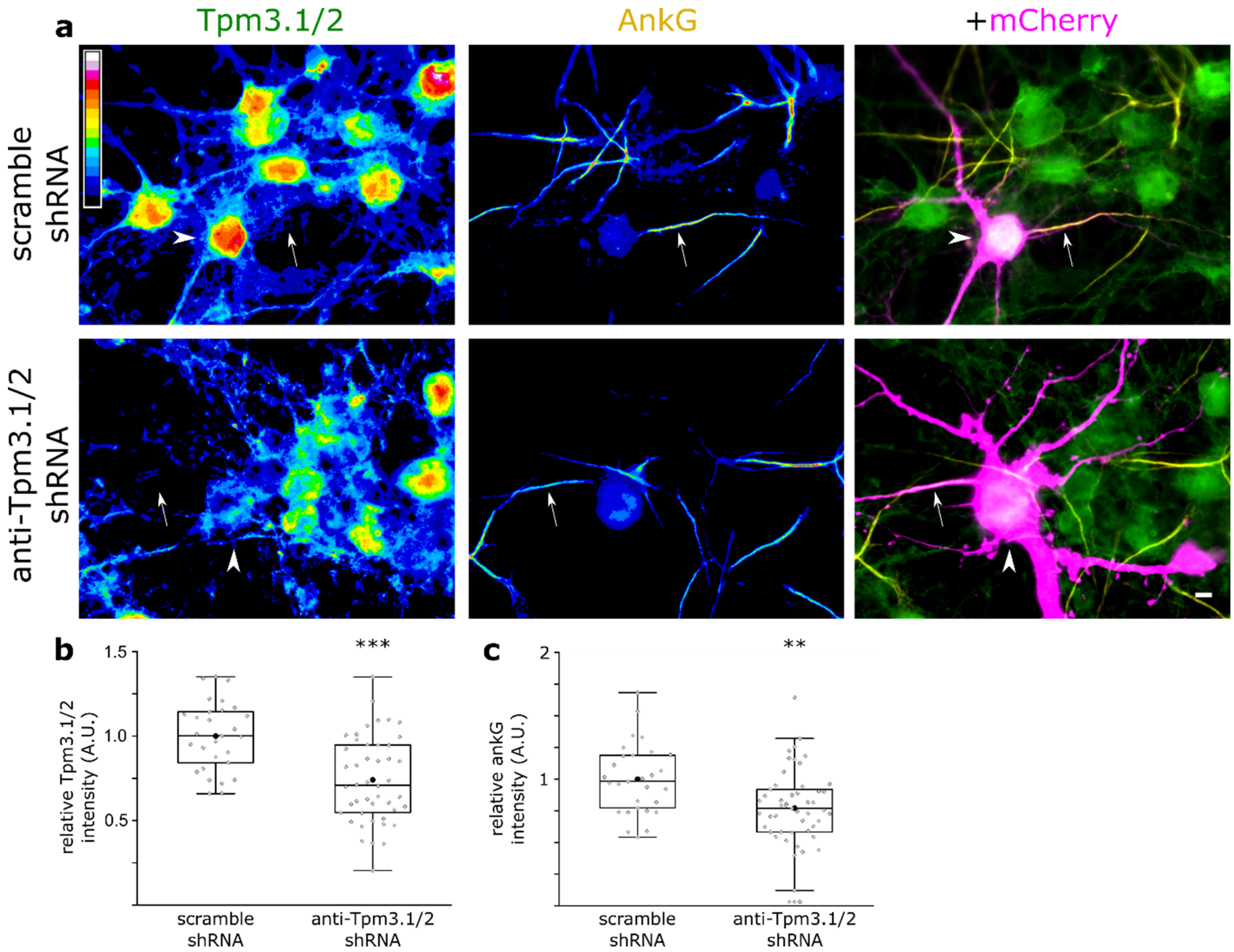


**Supplementary figure 11 | Tpm3.1/2 knockdown reduces the accumulation of ankyrin G at the AIS.** Rat hippocampal neurons transfected at 10 DIV using either anti-Tpm3.1/2 or scramble shRNA. Neurons were fixed at 14 DIV and stained using anti-γ/9d and anti-ankyrin G. **a** Anti-γ/9d and anti-ankyrin G immunofluorescence in neurons expressing scramble or anti-Tpm3.1/2 shRNA, and neighboring, non-transfected neurons. Arrowheads point to the transfected neuron, arrows point to axons. Color code indicates normalized fluorescence intensity levels. **b** Normalized relative intensity of anti-γ/9d fluorescence in the somata of neurons expressing scramble or anti-Tpm3.1/2 shRNA (scramble shRNA: 1 ± 0.04, mean ± SEM; shRNA: 0.74 ± 0.04, mean ± SEM, $p < 0.001$, *t* test). **c** Relative anti-ankyrin G fluorescence intensity in in each group (scramble shRNA: 1 ± 0.05, mean ± SEM; shRNA: 0.77 ± 0.04, mean ± SEM, $p < 0.01$, *t* test). Black circles represent mean value. Box borders represent the 25$^{th}$ and 75$^{th}$ percentiles, whiskers represent minimum and maximum values less than 1.5x the interquartile range lower or higher than the 25$^{th}$ or 75$^{th}$ percentiles, respectively (Tukey style). Scramble shRNA: n = 29, 3 independent experiments; anti-Tpm3.1/2 shRNA: n = 51, 3 independent experiments. * denotes statistical significance. **: $p < 0.01$; ***: $p < 0.001$. Scale bar: 5 μm.

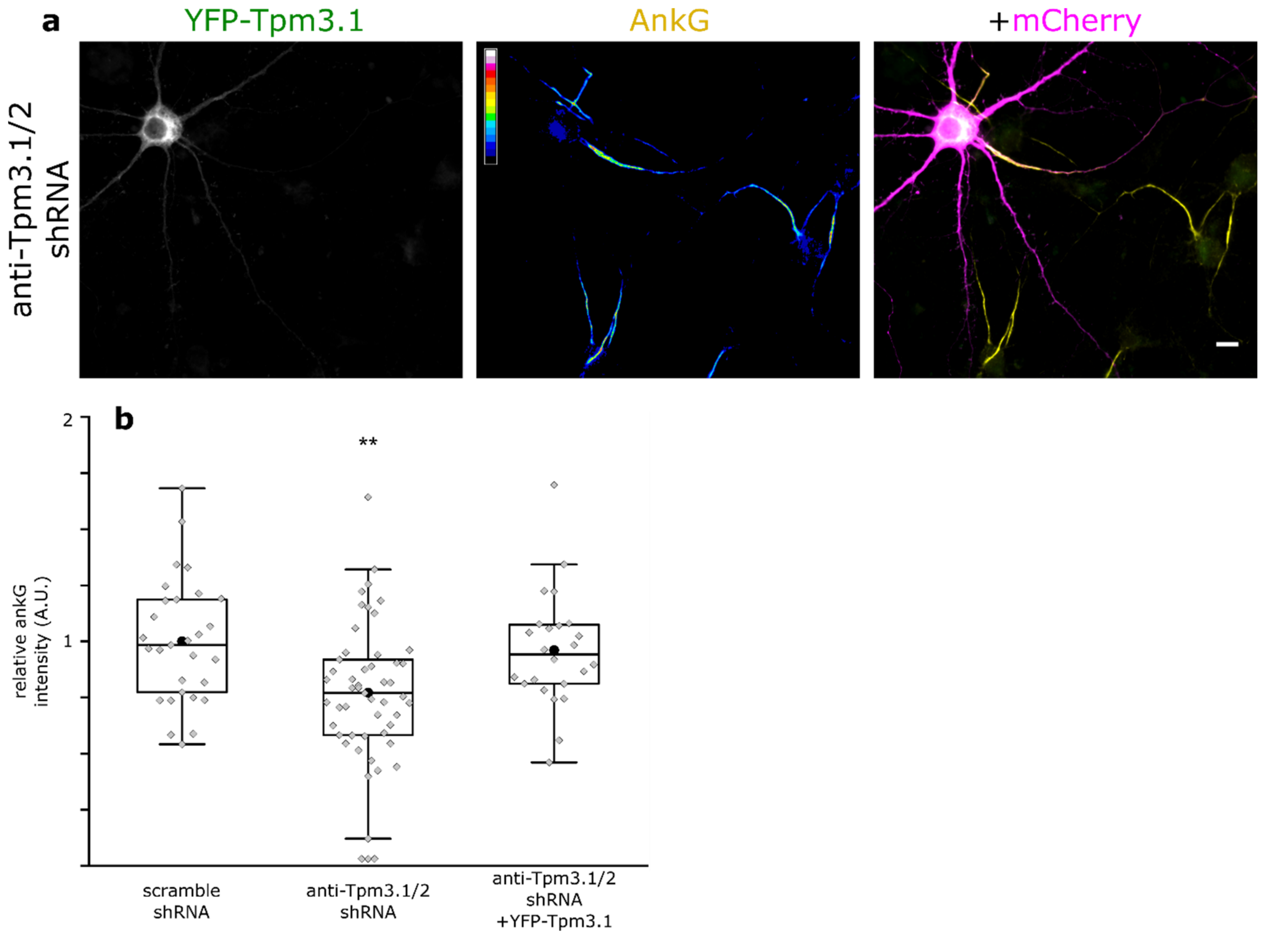


**Supplementary figure 12 | The effect of Tpm3.1/2 knockdown on the accumulation of ankyrin G at the AIS was rescued by the exogenous expression of human Tpm3.1.** Rat hippocampal neurons expressing mCherry-tagged anti-Tpm3.1/2 shRNA and YFP-tagged human Tpm3.1 at 10-14 DIV. Human Tpm3.1 differs from rat Tpm3.1 in the sequence within the targeted region. Anti-ankyrin G served to visualize ankyrin G. **a** Neuron expressing YFP-Tpm3.1 and anti-Tpm3.1/2 shRNA showing typical ankyrin G accumulation at the AIS. Color code indicates normalized fluorescence intensity levels. **b** Relative anti-ankyrin G fluorescence intensity in in each group (scramble shRNA: 1 ± 0.05, mean ± SEM; shRNA: 0.77 ± 0.04, mean ± SEM; anti-Tpm3.1/2 shRNA + YFP-Tpm3.1: 0.96 ± 0.05, mean ± SEM; $p < 0.01$, ANOVA, Tukey's test). Black circles represent mean value. Box borders represent the 25$^{th}$ and 75$^{th}$ percentiles, whiskers represent minimum and maximum values less than 1.5x the interquartile range lower or higher than the 25$^{th}$ or 75$^{th}$ percentiles, respectively (Tukey style). Anti-Tpm3.1 shRNA: n = 42, 2 independent experiments, partially used for Supplementary figure 11; scramble shRNA: n = 28, 2 independent experiments, partially used for Supplementary figure 11; anti-Tpm3.1/2 shRNA + YFP-Tpm3.1: n = 26, 2 independent experiments. * denotes statistical significance. **: $p < 0.01$. Scale bar: 5 μm.

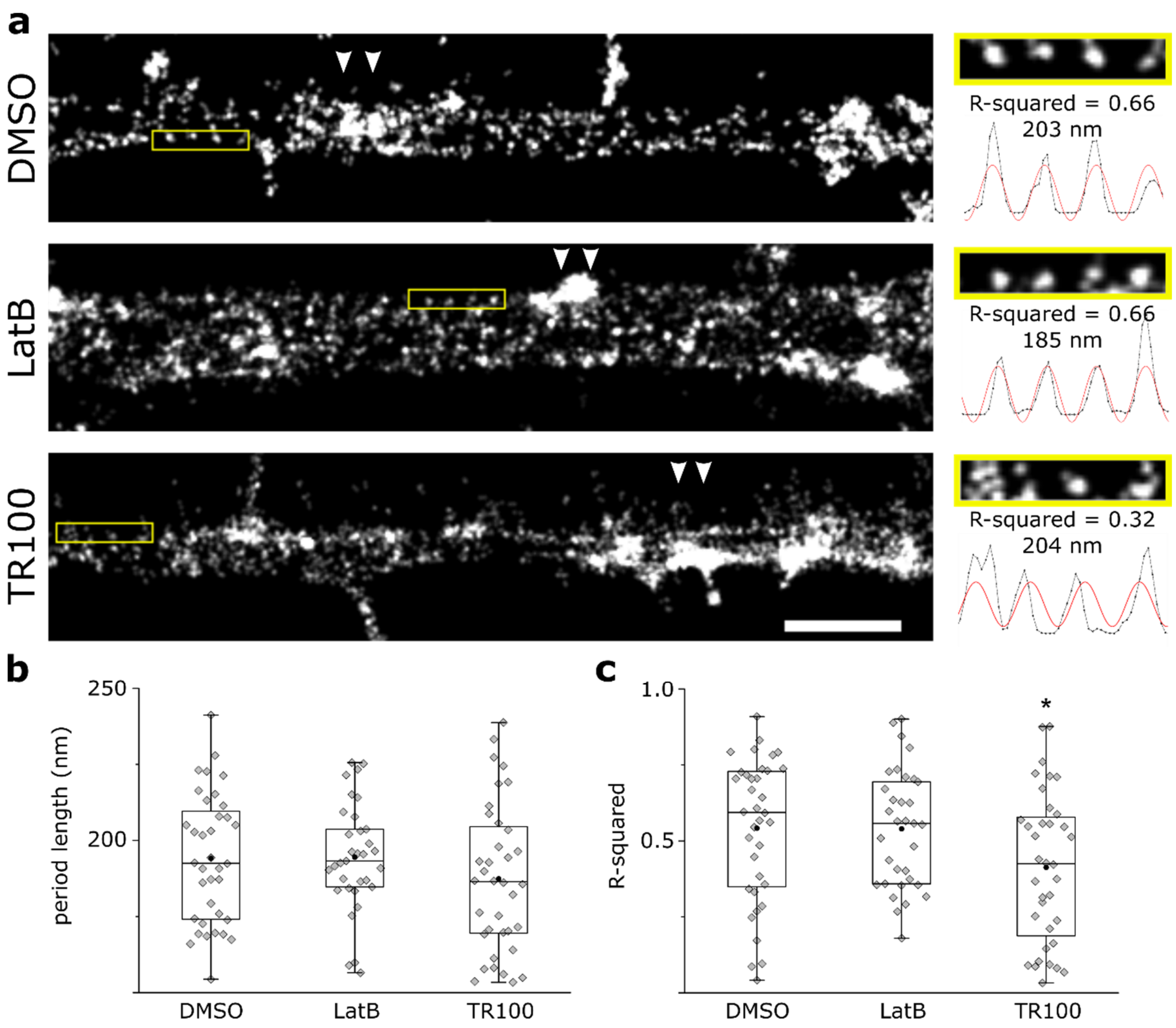


**Supplementary figure 13 | The periodicity of actin rings in the AIS is disrupted upon Tpm3.1 inhibition, but not latrunculin-B treatment. a** *Left*: STORM reconstructions of the AIS of rat hippocampal neurons at 14 DIV, treated overnight using either DMSO (0.2%), LatB (5 µM), or TR100 (5 µM). Alexa 647-tagged phalloidin served to label F-actin. Arrowheads point to AIS actin patches. *Right*: Higher magnification of the yellow-boxed area. The intensity profiles (black lines) and the best Fourier series fit (red lines) are shown. The period lengths and the R-squared values of the fits are indicated. 3 profiles were plotted in every neuron where periodicity was best visible. **b** Period lengths in each group. The difference in mean period length between groups was not statistically significant (ANOVA) **c** R-squared values of individual Fourier series fits in each group. TR100-treated neurons showed a significantly lower R-squared value, indicating a poorer Fourier series fit. LatB-treated neurons were not different from DMSO controls (Mann-Whitney *U* test). Black circles represent mean value. Box borders represent the 25$^{th}$ and 75$^{th}$ percentiles, whiskers represent minimum and maximum values less than 1.5x the interquartile range lower or higher than the 25$^{th}$ or 75$^{th}$ percentiles, respectively (Tukey style). DMSO: n = 12 neurons, 5 independent experiments; LatB: n = 11 neurons, 4 independent experiments; TR100: n = 12 neurons, 4 independent experiments. * denotes statistical significance. *: $p < 0.05$. Scale bar: 1 μm.
